# Kinase interplay centered on two regulatory elements compartmentalizes the checkpoint protein Lte1 in *S. cerevisiae*

**DOI:** 10.1101/2025.02.20.639281

**Authors:** Marco Geymonat, Qiuran Peng, Hanlu Chen, Sneha Parmar, Helen Mott, Darerca Owen, Marisa Segal

## Abstract

In budding yeast, Lte1, a component of the spindle position checkpoint, must be confined to the bud for correctly coupling chromosomal segregation across the bud neck with mitotic exit. Bud compartmentalization involves a phospho-dependent program with Ras-GTP acting as the ultimate cortical anchor. Here we have dissected this program by identifying an Lte1 domain participating in Ras binding and two phospho-regulatory motifs, the N- and C-LREs (Lte1 Regulatory Elements). The C-LRE mediates control of Lte1 cortical recruitment by the p21-activated kinase, Cla4. The N-LRE modulates Lte1 binding to Ras more directly and is phosphorylated by the yeast MARK/PAR-1 protein kinases, Kin1/2. Along with previously described CDK phosphorylation sites, the N- and C-LREs are required for Lte1 localization and timely mitotic exit. These studies uncover the coordinated interplay between multiple protein kinases allowing Lte1 to integrate cell polarity and cell cycle cues for strict compartmentalization at the bud cell cortex.

## INTRODUCTION

In asymmetric cell divisions, the cell polarity and cell cycle machineries collaborate to define opposite cortical compartments with the intended division site in between. These not only support navigational cues to bias the orientation of the mitotic apparatus, but also lay out the foundations for differential daughter cell identity and age (Casas Gimeno and Paridaen, 2022; Guo and Dong, 2022; Habib and Acebron, 2022; Pineau et al., 2023; Yamashita, 2023).

Mitotic exit control in the budding yeast *Saccharomyces cerevisiae* is a remarkable paradigm for exploring how cortical domain demarcation underpins coordination between the mitotic apparatus and cytokinesis (Baro et al., 2017; Caydasi and Pereira, 2012; Geymonat and Segal, 2017). Highly polarized growth in yeast generates a daughter (bud) connected to the mother cell through the bud neck, the future cell division site (Chiou et al., 2017; Marquardt et al., 2021; Okada et al., 2013; Spiliotis and McMurray, 2020). Crucially, mitotic exit can only occur after the mitotic spindle elongates across the bud neck, thus bringing a set of the duplicated chromosomes into the bud. This precondition hinges on the regulated release of the phosphatase Cdc14 from sequestration in the nucleolus by two pathways acting in tandem: the Fourteen Early Anaphase Release (FEAR) network and the Mitotic Exit Network (MEN). Upon sustained release of Cdc14 by the MEN (Queralt and Uhlmann, 2008; Rock and Amon, 2009; Shou et al., 1999; Stegmeier et al., 2002; Visintin et al., 1999), dephosphorylation of critical cyclin-dependent kinase (CDK) substrates, sets in motion irreversible mitotic exit and cytokinesis (Jaspersen et al., 1999; Jaspersen et al., 1998; Visintin et al., 1998).

The Spindle Position Checkpoint (SPOC) restricts MEN activation until one pole of the spindle has entered the bud (Bardin et al., 2000; Caydasi and Pereira, 2012; Pereira et al., 2000). The SPOC controls the small GTPase Tem1 (Scarfone and Piatti, 2015; Shirayama et al., 1994b), the upstream trigger of the MEN protein kinase cascade. Tem1 is localized to the Spindle Pole Body (SPB, analog of the animal centrosome) along with its negative regulator Bub2-Bfa1, a two-component GTPase-activating protein (GAP) that stimulates conversion of GTP-bound Tem1 to its GDP-bound state (Bardin et al., 2000; Geymonat et al., 2002; Pereira et al., 2000; Shirayama et al., 1994b). To date, a Tem1 guanine nucleotide exchange factor (GEF) has not been identified. Instead, two protein kinases modulate Tem1 via its GAP in an antagonistic fashion. Cdc5 inhibitory phosphorylation of Bfa1 results in active Tem1 (Geymonat et al., 2003; Hu et al., 2001). Conversely, Kin4 blocks GAP downregulation by Cdc5 keeping Tem1 inactive. Importantly, Kin4 activity is confined to the mother cell (D’Aquino et al., 2005; Pereira and Schiebel, 2005) and opposed by Lte1, a protein restricted to the bud cortex (Bardin et al., 2000; Pereira et al., 2000).

Lte1 shares sequence similarity with the catalytic module of Ras-GEFs (Boguski and McCormick, 1993; Shirayama et al., 1994a). In canonical Ras-GEFs, this module consists of two elements (Bos et al., 2007): the Ras Exchanger Motif (REM) and the Cdc25 Homology Domain (CHD) that constitutes the GEF active site (Broek et al., 1987; Camonis et al., 1986; Robinson et al., 1987). In Lte1, separate REM and CHD-like sequences map to Lte1’s N and C-termini, respectively and have been implicated in Lte1 cortical anchoring via Ras-GTP (Geymonat et al., 2009; Geymonat et al., 2010; Seshan and Amon, 2005; Yoshida et al., 2003), while a central region of Lte1 binds and inhibits Kin4 (Bertazzi et al., 2011; Falk et al., 2016a; Falk et al., 2011).

These findings laid the premise for the "two-zone model" in which the SPB supports a spatial sensor, the Bub2-Bfa1-Tem1 module, responding to transit across the bud neck (Bertazzi et al., 2011; Chan and Amon, 2010; Falk et al., 2011). Briefly, the sensor becomes polarized to the SPB committed to the bud during spindle alignment (Caydasi and Pereira, 2009; Monje-Casas and Amon, 2009; Pereira et al., 2001). Upon elongation of the spindle across the bud neck, Tem1 on the leading SPB escapes Kin4 inhibitory action by entering the Lte1-marked bud compartment in which GAP downregulation by Cdc5 and concomitant Tem1 activation take effect (Caydasi and Pereira, 2009; Falk et al., 2016b; Fraschini et al., 2006; Monje-Casas and Amon, 2009). Accordingly, ectopic Lte1 localization to the mother cell abrogates the SPOC (Bertazzi et al., 2011). However, the role of Tem1 as a canonical GTPase switch has been brought to question in a study positing instead that the GTPase cycle would govern Tem1 effective concentration at the leading SPB for activation of the MEN kinase cascade (Zhou et al., 2024). Furthermore, Lte1 may additionally activate mitotic exit in a Kin4-independent manner by an, as yet, unknown mechanism (Caydasi et al., 2017; Falk et al., 2016a).

Cdc42, the master regulator of cell polarity (Chiou et al., 2017), is required for Lte1 cortical recruitment at the incipient bud following G1-CDK activation. Yet, it is dispensable for maintaining Lte1 bud cortex association, suggesting multiple mechanisms are at play during bud growth. An intact septin ring is also essential to confine Lte1 to the bud (Hofken and Schiebel, 2002; Jensen et al., 2002; Seshan et al., 2002).

Lte1 exhibits prominent cell cycle-dependent phosphorylation (Bardin et al., 2000; Lee et al., 2001) attributed to two protein kinases: the p21-activated kinase (PAK) Cla4 and CDK (Bardin et al., 2000; Hofken and Schiebel, 2002; Jensen et al., 2002; Seshan et al., 2002; Ubersax et al., 2003). Lte1 phosphorylated state, its cortical association and binding to Ras may be linked (Geymonat et al., 2010; Jensen et al., 2002; Seshan and Amon, 2005; Seshan et al., 2002) and are reversed after Cdc14 release (Jensen et al., 2002; Seshan and Amon, 2005; Seshan et al., 2002). Ras and Cla4 may be interdependent for promoting Lte1 phosphorylation and establishing Lte1-Ras complexes at the cell cortex (Seshan and Amon, 2005). Accordingly, Lte1 bulk phosphorylation and cortical association are abrogated in either *cla4Δ* or *ras1Δ ras2Δ* mutants (Hofken and Schiebel, 2002; Seshan and Amon, 2005; Yoshida et al., 2003).

Here we have dissected phospho-dependent control of Lte1 cortical recruitment and compartmentalization with reference to Lte1-Ras complex formation. First, we defined the minimal Lte1 sequence requirements for this program. This analysis implicated the REM in Lte1 cortical anchoring via Ras-GTP and identified two distinct phospho-site clusters as Lte1 Regulatory Elements or LREs. The C-LRE next to the CHD was a target of Cla4 and a phospho-mimetic substitution at a PAK consensus site in this element restored correct Lte1 localization in a *cla4Δ* background. By contrast, phospho-mimetic substitutions in the N-LRE near the REM enhanced intrinsic binding to Ras-GTP resulting in Lte1 breaching compartmentalization with the concomitant abrogation of the SPOC. Furthermore, we found that the yeast MARK/PAR-1 protein kinases Kin1/2 (Levin et al., 1987; Wu and Griffin, 2017; Yuan et al., 2016) contributed to N-LRE phosphorylation and ensuing enhancement of Lte1 binding to Ras-GTP. Finally, cancelling phospho-sites in the N-LRE and C-LRE along with previously described CDK sites (Jensen et al., 2002) disrupted both Lte1 localization program and timely mitotic exit. Taken together, our studies uncover an orderly protein kinase interplay allowing Lte1 to integrate cell polarity and cell cycle controls enforcing its strict compartmentalization at the bud cell cortex.

## RESULTS

### Exploring Cla4-dependent control of Lte1 localization by still-imaging analysis

Consistent with previous studies (Bardin et al., 2000; Hofken and Schiebel, 2002; Seshan et al., 2002), the mobility shift of a Lte1-Protein A (ProA) fusion due to phosphorylation *in vivo* was markedly suppressed in a *cla4Δ* mutant (Figure 1 A). Yet, according to the *Saccharomyces* Genome Database (SGD, https://www.yeastgenome.org/), Lte1 undergoes phosphorylation at ∼80 sites, with only 2 responding to the PAK consensus R/K-R-X-pS/T and 4 to the minimal Cla4 consensus R-X-pS/T (Mok et al., 2010). Lte1 is a known CDK substrate (Jensen et al., 2002) but other unidentified protein kinases must target Lte1.

**Figure 1.**
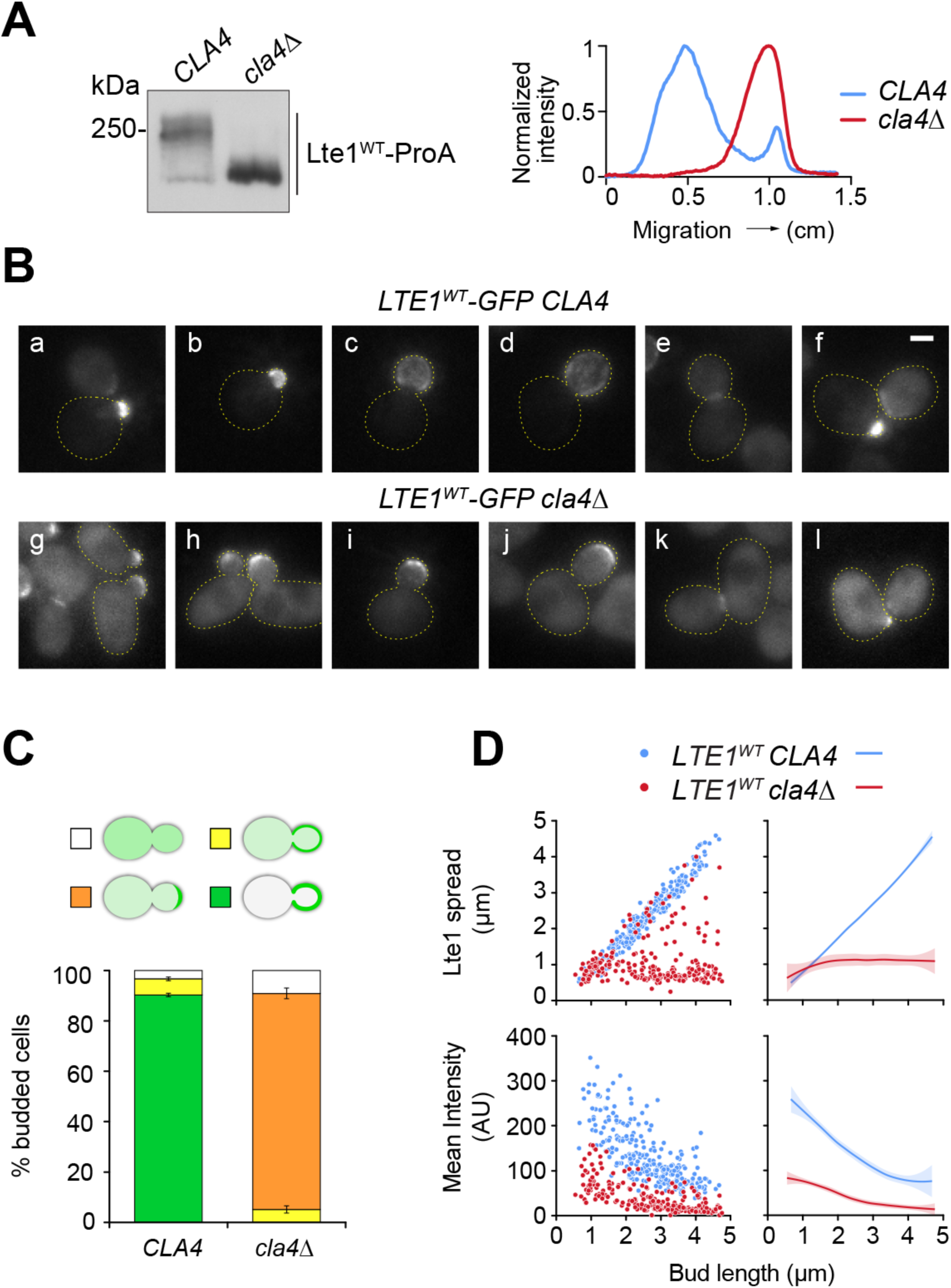
Program of Lte1 cortical localization and its dependence on the p21-activated kinase Cla4. (A) Contribution of Cla4 to bulk phosphorylation of Lte1 assessed by western blot analysis of whole extracts of wild type vs *cla4Δ* cells expressing Lte1 fused to Protein A (ProA), after arrest in metaphase (left panel). Mobility shift was evaluated using intensity linescans along the direction of electrophoretic migration (right panel). Inactivation of Cla4 virtually abrogated the phospho-dependent mobility shift as previously reported (Hofken and Schiebel, 2002; Seshan et al., 2002). (B) Fluorescence images selected from asynchronous populations representing stages for Lte1-GFP localization along the cell cycle in wild type vs *cla4Δ* cells. Scale bar, 2 µm. Expression levels of Lte1-GFP in the strains analyzed are presented in Suppl. Figure 1 B. (C) Modes of Lte1 localization in wild type vs *cla4Δ* budded cells scored collectively in still images from asynchronous populations. The plot presents the mean of 3 independent counts of 120 cells ± SEM. The categories scored are illustrated in Suppl. Figure 1 C. (D) Scatterplot (left) and smoothed conditional mean plot (right) for single-cell analysis of Lte1-GFP cortical spread along the bud cell polarity axis (the width of the linescan peak at half maximal value) and mean intensity in the bud as a function of bud length in wild type (n = 258) vs *cla4Δ* (n = 223) cells, derived from fluorescence intensity linescans (see Materials and Methods). Solid lines correspond to the smoothed conditional mean (locally weighted scatterplot smoothing) and shading represents the 95% CI. Cumulative distribution of bud length in each sample is shown in Suppl. Figure 1 D.

In order to genetically dissect the phosphoregulatory mechanisms underlying the Lte1 program of localization, a quantitative framework based on live imaging was developed and applied, in the first instance, to otherwise wild type or *cla4Δ* cells expressing Lte1^WT^-GFP (Figure 1 B-D). Images of cells from asynchronous populations ordered by morphology recapitulated the wild type program of Lte1 cortical localization in agreement with time lapse data (Suppl. Figure 1 A). Briefly, Lte1^WT^-GFP began marking the incipient bud (Figure 1 B, a) and extended over the growing bud cortex (Figure 1 B, b-d). After mitotic exit, label was delocalized, transiently marking the division site (Figure 1 B, e). Finally, Lte1-GFP in the former mother cell was recruited to a new budding site while forming a broad crescent at the recent division site in the daughter cell (Figure 1 B, f) until a new bud site was selected. By contrast, Lte1^WT^-GFP was only weakly recruited to the incipient bud over cytoplasmic label background in *cla4Δ* cells (Figure 1 B, g), stayed apical during bud growth (Figure 1 B, h-j) and returned to the division site at mitotic exit (Figure 1 B, k) until redirected to the new bud site (Figure 1 B, l). The mutation did not affect Lte1^WT^-GFP expression level (Suppl. Figure 1 B). Weak apical localization was the predominant mode of label collectively scored in *cla4Δ* budded cells (Figure 1 C and Suppl. Figure 1 C).

Single-cell linescan analysis for fluorescence intensity along the mother-bud axis was implemented to obtain a one-dimensional quantitative description of Lte1-GFP distribution in terms of cortical spread or mean intensity as a function of bud cell length (Figure 1 D). Lte1 spread correlated with bud length throughout in wild type cells, while mean intensity peaked in small-budded cells and declined with bud growth. A similar decline was apparent when measuring mean Lte1-GFP cortical intensity in the entire bud in a time lapse series (Suppl. Figure 1 A). For validation, average linescan traces at 0.5 µm-bud length intervals were derived from the original dataset (Suppl. Figure 1 D and E). In contrast to wild type cells, spread of Lte1^WT^ in *cla4Δ* cells mostly failed to scale with bud growth while initial mean intensity at the bud cell cortex was reduced and further declined with bud growth. Average linescan traces from the *cla4Δ* dataset (Suppl. Figure 1 E) reflected both weak apical localization and increased cytoplasmic background.

Taken together, a *cla4Δ* mutation virtually eliminated Lte1 phosphorylation and reduced Lte1 cortical association down to limited apical label throughout bud growth. This subtle localization retained by the *cla4Δ* strain has not been previously reported.

### Lte1 sequences required for its phospho-dependent cortical association

In the human Ras-GEF Son of Sevenless 1 (hSOS1), binding of Ras-GTP to the REM allosteric site enhances GDP-GTP exchange by the CHD (Margarit et al., 2003; Sondermann et al., 2004). In Lte1, the REM and CHD sequences are required for efficient Lte1 cortical association (Jensen et al., 2002; Yoshida et al., 2003). In order to narrow down Lte1 sequences critical for phospho-dependent control, in-frame internal deletions preserving the REM and CHD were generated (Figure 2 A). A minimal Lte1 version lacking amino acids 250 to 950 retained cortical localization still dependent on Cla4 (Figure 2 B and C).

**Figure 2.**
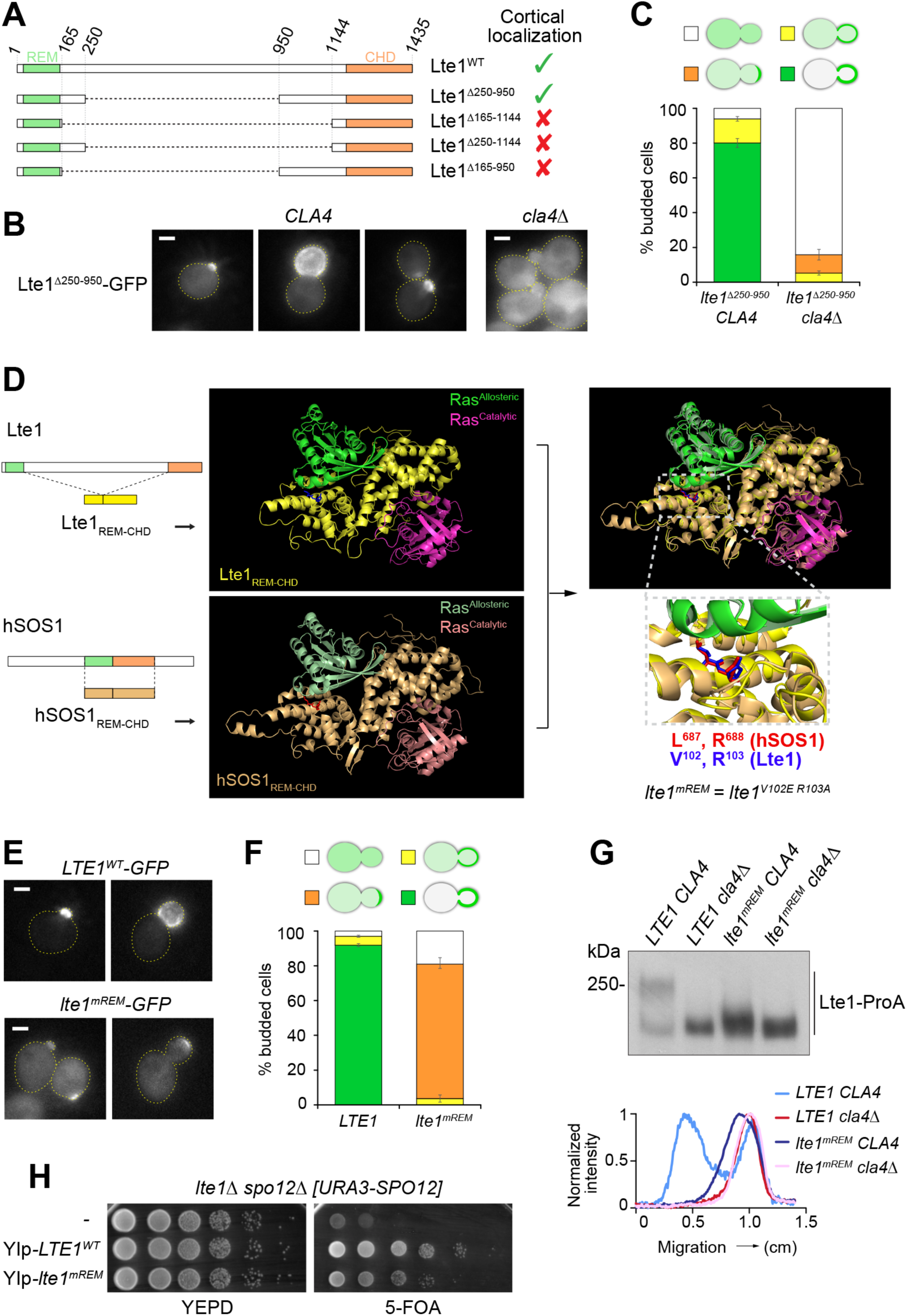
Lte1 sequence requirements for phospho-dependent cortical localization. (A) Minimal construct retaining Lte1 cortical localization. Sequences in Lte1^Δ250-950^ adjacent to the REM (green box) and CHD (orange box), respectively, were required. (B) Representative images for Lte1^Δ250-950^-GFP expressed in wild type vs *cla4Δ* cells. Lte1^Δ250-950^ retained cortical localization dependent on Cla4. Scale bar, 2 µm. (C) Distribution of modes of Lte1^Δ250-950^-GFP cortical localization in budded cells (as defined in Suppl. Figure 1 C) in wild type or *cla4Δ* mutant backgrounds. The plot shows the mean ± SEM of three independent counts of 120 cells. (D) Strategy for identifying critical residues in the putative REM of Lte1 that may be required for interaction with Ras-GTP. Modeled structure of an Lte1_REM-CHD_ unit based upon hSOS1_REM-CHD_ pinpointed V^102^ and R^103^ (blue) as counterparts of key residues in hSOS1 (red) required for Ras-GTP ("Ras allosteric") interaction with the REM (Sondermann et al., 2004). The mutant encoding V102E and R103A substitutions was referred to as *lte1^mREM^*. (E) Representative fluorescence images for localization of Lte1^WT^-GFP vs Lte1^mREM^-GFP. Cortical localization was reduced to subtle apical retention by substitutions within the putative REM. Scale bar, 2 µm. (F) Distribution of modes of Lte1^WT^ vs Lte1^mREM^ cortical localization in budded cells. The plot shows the mean ± SEM of three independent counts of 120 cells. (G) Western blot analysis of whole cell extracts of the indicated strains following synchronization at metaphase (upper panel), paired with intensity linescans along the direction of electrophoretic migration (lower panel). Bulk phosphorylation of Lte1^mREM^ was markedly decreased and further reduced in a *cla4Δ* background. (H) Impaired suppression of *lte1Δ spo12Δ* synthetic lethality by *lte1^mREM^*. *lte1Δ spo12Δ [URA3-SPO12]* cells transformed with integrative plasmids encoding *LTE1* or *lte1^mREM^* were spotted onto YEPD or 5-FOA plates to assess the ability of the integrative constructs to support viability upon counter-selection of the resident *URA3-SPO12* autonomous plasmid.

Unlike canonical Ras-GEFs, Lte1 binds solely to Ras-GTP *in vitro* and *in vivo* (Seshan and Amon, 2005; Yoshida et al., 2003). Furthermore, human H-Ras can replace yeast Ras proteins as the Lte1 cortical anchor (Suppl. Figure 2 A-B). These observations support the idea of functional conservation, providing a rationale for investigating the direct involvement of the REM. Accordingly, the REM and CHD domains of Lte1 were modelled with two H-Ras molecules bound, using the structure of the hSOS1 complex (PDB 1nvv) as a template (Figure 2 D). Based on this model, substitutions V^102^ to E and R^103^ to A were deemed analogous to those used previously to prevent Ras-GTP binding to the hSOS1 REM (L687 to E and R688 to A; Sondermann et al., 2004). An allele encoding these substitutions is designated *lte1^mREM^*.

Lte1^mREM^-GFP was poorly recruited at the incipient bud and weakly labeled the apical cortex of midsize buds in otherwise wild type cells (Figure 2 E and F). This label was too weak for digital quantification. Localization resembled that seen when Lte1^WT^-GFP was expressed in cells lacking yeast *RAS* genes, supporting the idea that substitutions within the REM disabled anchorage via Ras (Suppl. Figure 2 A-B). Furthermore, western blot analysis indicated a marked decrease in bulk phosphorylation of Lte1^mREM^-ProA *in vivo*, with a subtle mobility shift dependent on Cla4 retained (Figure 2 G). Finally, substitutions within the REM undermined Lte1’s ability to rescue the synthetic lethality between *lte1Δ* and the FEAR mutation *spo12Δ* (Figure 2 H).

In conclusion, sequences adjacent to the REM and CHD might contain phospho-regulatory sites supporting Cla4-dependent Lte1 localization. In addition, the Lte1 REM might act as a Ras-GTP binding site analogous to that in hSOS1 and be key to Ras-GTP mediated cortical association.

### Phospho-mimetic substitutions at sites adjacent to the REM directed Lte1 to the mother cell cortex, independent of Cla4

Previous phospho-proteomic studies (Holt et al., 2009; Lanz et al., 2021) identified a 6 phospho-site cluster proximal to the REM, retained in Lte^Δ250-950^ (Figure 3 A). To probe its significance, we reasoned that phospho-mimetic substitutions would be particularly revealing, given the correlation between Lte1 phosphorylated state and cortical association. In view of our results, we now refer to this motif as N-Lte1 Regulatory Element or N-LRE.

**Figure 3.**
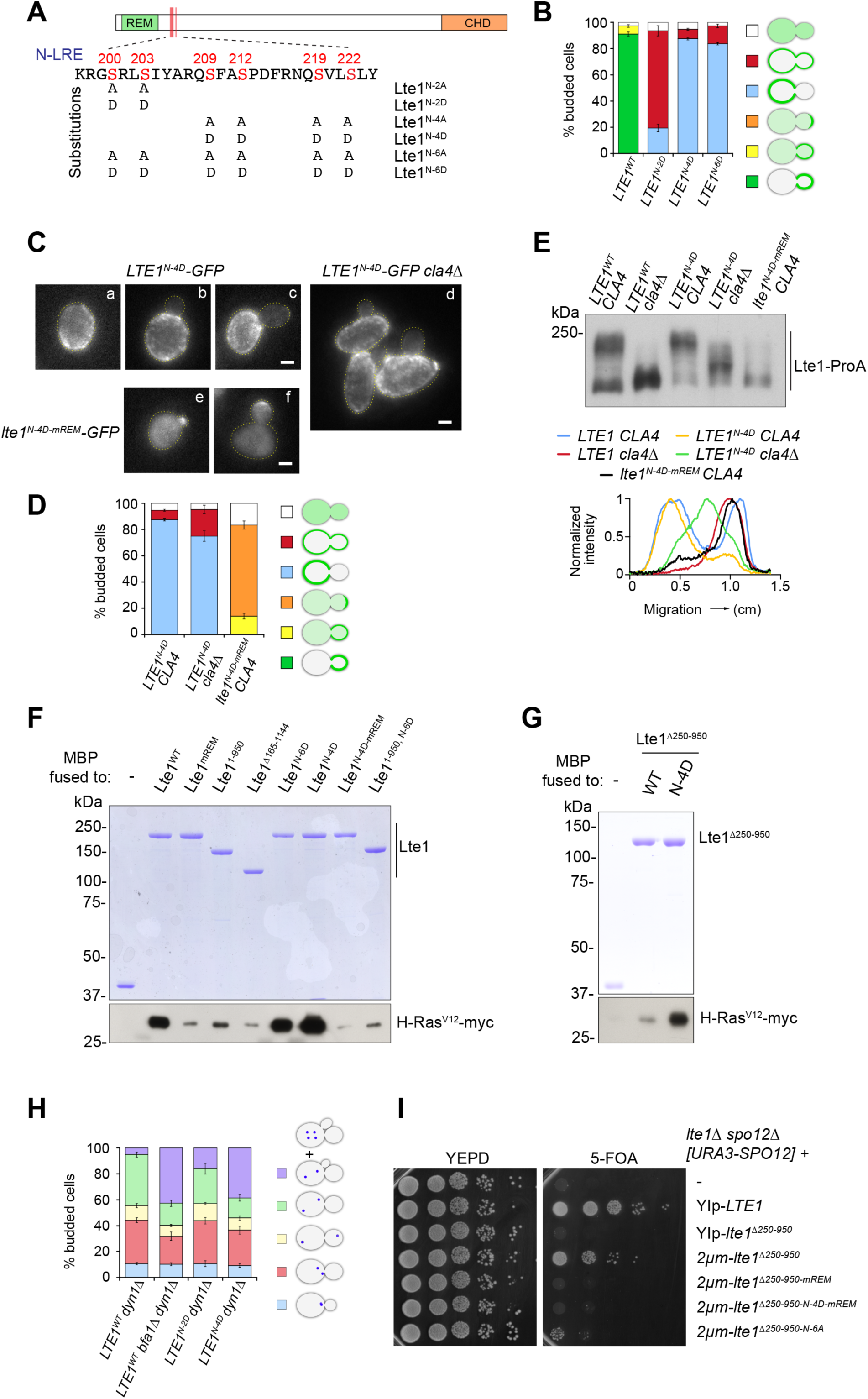
A cluster of phospho-sites proximal to the REM constitutes a regulatory element of Lte1’s cortical program. (A) Diagram outlining the position and sequence of the N-terminal Lte1 regulatory element (N-LRE). Phospho-sites as curated by the SGD and selected for genetic analysis are shown in red. Also shown is a summary of substitutions created and product designation for the purpose of this study. (B) Phospho-mimetic substitutions directed Lte1 to the mother cell cortex, thus breaching Lte1 compartmentalization. Distribution of modes of Lte1-GFP localization scored in budded cells included two new categories — mother or mother and bud cell cortex (as illustrated in Suppl. Figure 2 C). The plot corresponds to mean ± SEM of three independent counts of at least 120 cells. (C) Localization of Lte1^N-4D^ to the mother cell cortex (a-unbudded, b-small budded and c-large budded cells) persisted in a *cla4Δ* mutant background (d) but was abrogated when further substitutions at the REM were introduced (e and f). Scale bar, 2 µm. (D) Distribution of modes of localization in budded cells of the indicated strains (according to the categories outlined in Suppl. Figure 2 C) showing mean ± SEM of three independent counts of at least 120 cells. (E) Western blot (upper panel) and intensity linescan analyses (lower panel) for Lte1 bulk phosphorylation assessed by mobility shift in whole extracts of cells expressing the indicated versions of Lte1-ProA in wild type or *cla4Δ* backgrounds, prepared after metaphase arrest. Phospho-mimetic substitutions in the N-LRE restored Lte1 mobility shift, dependent on an intact REM, in a *cla4Δ* background. (F) Phospho-mimetic substitutions in the N-LRE enhanced Lte1’s interaction with Ras-GTP in an *in vitro* pull-down assay. The indicated versions of Lte1 fused to MBP and H-Ras^V12^-myc (an oncogenic mutant version that is constitutively bound to GTP, see Materials and Methods) were purified from yeast. After resolving the pull-down product by SDS-PAGE the lower part of the gel was excised and subjected to western blot analysis to visualize bound Ras while the top part was directly stained by Coomassie blue to view control MBP or MBP-Lte1 fusion baits. (G) Phospho-mimetic substitutions in the N-LRE also enhanced H-Ras^V12^-myc binding to MBP-Lte1^Δ250-950^ *in vitro*. Analysis of the pull-down was performed as in (F). (H) Phospho-mimetic substitutions within the N-LRE that disrupted Lte1 compartmentalization impaired the SPOC. Budded cells from cultures incubated at 14°C were scored using Spc29-CFP to visualize SPBs, according to the categories depicted in the cartoons (blue dots represent SPBs). In addition to cells containing unseparated SPBs (light blue), short spindles (red) and elongated spindles across the bud neck (yellow), a *dyn1Δ* mutation increased spindle misorientation, causing otherwise wild type cells to accumulate large budded cells with mispositioned anaphase spindles (green) in response to the SPOC. A *bfa1Δ* mutation abrogates the SPOC allowing cells to exit mitosis despite the presence of both spindle poles within the mother cell. This translates into an increase in cells undergoing re-budding and SPB re-duplication (violet). This latter category was also increased in *LTE1^N-2D^* and *LTE1^N-4D^*strains. The plot depicts the mean ± SEM of 3 independent counts of at least 120 cells. Representative images for each category scored are shown in Suppl. Figure 2 G. (I) Substitutions that cancel phosphorylation at the N-LRE abrogated rescue of *lte1Δ spo12Δ* synthetic lethality in the context of multicopy *lte1^Δ250-950^*. *lte1Δ spo12Δ [URA3-SPO12]* cells transformed with the indicated integrative (YIp) or multicopy (2µm) plasmids were spotted onto YEPD or 5-FOA plates to assess viability upon counter-selection of the resident *URA3-SPO12* autonomous plasmid.

Otherwise wild type strains expressing the mutant versions Lte1^N-2D^, Lte1^N-4D^ or Lte1^N-6D^ (Figure 3 A) fused to GFP exhibited label at the mother cell cortex, demonstrating a breach in compartmentalization. Lte1^N-4D^ and Lte1^N-6D^ strongly favored the mother cell (Figure 3 B and Suppl. Figure 2 C).

Further characterization was pursued with Lte1^N-4D^ (Figure 3 C-E). Deregulated cortical association of Lte1^N-4D^-GFP was apparent in 92 % of unbudded cells (Figure 3 C, a) and favored mother cells past bud emergence (Figure 3 C, b-c and Figure 3 D). Lte1^N-4D^ cortical association was independent of Cla4 (Figure 3 C, d and Figure 3 D). Yet, localization was markedly lost with the addition of REM substitutions presumed to disrupt binding to Ras-GTP (Lte1^N-4D-mREM^, Figure 3 C, e-f and Figure 3 D). Thus, phospho-mimetic residues could bypass phospho-regulation but not the requirement for an intact REM.

In view of the close relationship between Lte1 cortical association, phosphorylation and complex formation with Ras-GTP (Bardin et al., 2000; Geymonat et al., 2010; Seshan and Amon, 2005; Seshan et al., 2002), the impact of phospho-mimetic substitutions in the N-LRE on those processes was explored.

First, bulk phosphorylation of Lte1 *in vivo* was assessed in wild type or *cla4Δ* strains expressing Lte1^WT^ or Lte1^N-4D^ fused to ProA (Figure 3 E). In contrast to Lte1^WT^, Lte1^N-4D^ retained a considerable mobility shift in a *cla4Δ* background (as was the case for Lte1^N-2D^ and Lte1^N-6D^, see Suppl. Figure 2 D). Yet, the mobility shift was suppressed by the REM substitutions that impaired cortical association (Figure 3 E). These data suggest that Cla4-independent cortical association restored Lte1 phosphorylation by other protein kinases outside the N-LRE.

Second, Lte1 binding to Ras-GTP *in vitro* was assessed by pull-down assays using purified Lte1^WT^ or mutants fused to MBP as bait (Figure 3 F and G). Relative to Lte1^WT^, substitutions within the REM (Lte1^mREM^) or an in-frame deletion encoding solely joint REM-CHD sequences (Lte1^Δ165-1144^) markedly reduced binding to Ras-GTP while the N-terminal Lte1^1-950^ fragment retained limited binding (Figure 3 F). Phospho-mimetic substitutions in the N-LRE in full length Lte1 (Lte1^N-6D^ or Lte1^N-4D^, Figure 3 F) or in the context of Lte1^Δ250-950^ (Figure 3 G) enhanced binding to Ras-GTP, an effect abrogated by additional substitutions in the REM (Lte1^N-4D-mREM^, Figure 3 F). Yet, N-6D substitutions did not enhance binding of Lte1^1-950^, underscoring a requirement for a C-terminal domain. Pull-down assays followed by quantitative western blot analysis confirmed that N-2D, N-4D or N-6D substitutions increased Lte1 binding to Ras-GTP (Suppl. Figure 2 E-F). Taken together, phospho-mimetic substitutions within the N-LRE deregulated Lte1-Ras complex formation and cortical association, thus superseding cell cycle-dependent phosphoregulatory control.

Mutants impaired for spindle positioning across the bud neck, e.g., *dyn1Δ* (a deletion of the dynein heavy chain gene) accumulate budded cells with elongated spindles in the mother cell upon SPOC-enforced delay, a phenotype enhanced at 14°C (Li et al., 1993). According to the "two-zone model", Lte1 strict confinement to the bud underscores checkpoint proficiency. The impact of phospho-mimetic substitutions at the N-LRE on checkpoint performance was therefore tested in a *dyn1Δ* background by scoring collectively re-budding and SPB re-duplication to determine the proportion of cells undergoing inappropriate progression into the next cell cycle (Figure 3 H and Suppl. Figure 2 G).

As expected, the *LTE1^WT^ dyn1Δ* culture was enriched for large-budded cells with mispositioned anaphase spindles. Additionally introducing *bfa1Δ*, a mutation that disables the SPOC, depleted this category and increased cells undergoing re-budding and SPB re-duplication. Similarly, *LTE1^N-4D^ dyn1Δ* and, to a lesser extent *LTE1^N-2D^ dyn1Δ* strains pre-empted mitotic exit (Figure 3 H). It follows that phospho-mimetic substitutions within the N-LRE proved sufficient not only to breach Lte1 compartmentalization but also to disrupt the checkpoint, highlighting the impact of deregulated Lte1 binding to Ras.

The effect of impeding phosphorylation at the N-LRE was assessed in the simplified context of Lte1^Δ250-950^. *lte1^Δ250-950^* suppressed *lte1Δ spo12Δ* synthetic lethality when carried in a multicopy plasmid (Figure 3 I). In the same context, substitutions at the REM, even when combined with the phospho-mimetic N-4D prevented rescue. Finally, substitutions that cancelled phosphorylation at the N-LRE (N-6A) prevented suppression, showing that those phospho-sites are required for Lte1^Δ250-950^ function *in vivo*.

### C-LRE, a second phosphoregulatory site adjacent to the CHD

Lte1 is a substrate of Cla4 and CDK. To establish a functional link to particular phospho-sites, *in vitro* kinase assays were performed with purified Lte1 substrates designed to focus on the N-LRE (that contains at least one potential phosphorylation site for each kinase). Analogue-sensitive CDK-as complexes (Bishop et al., 2000), Cla4 and Cla4^KD^ (kinase-dead) were also purified from yeast (Figure 4 A-C). The N-terminal Lte1^1-325^ was a specific substrate for both CDK and Cla4 (Figure 4 B), as phosphorylation was prevented by the inhibitory analog 1NM-PP1 or Cla4^KD^, respectively. However, phosphorylation persisted despite introducing N-2A, N-4A or N-6A substitutions. An in-frame deletion consisting of the REM and CHD (Lte1^Δ165-1144^) showed reduced phosphorylation with either kinase. Using Lte1^Δ250-950^ as substrate, radioactive incorporation increased but substitutions within the N-LRE failed to reveal phosphorylation at that sequence by CDK or Cla4. Importantly, a deletion eliminating the sequence proximal to the CHD only (Lte1^Δ250-1144^) reduced phosphorylation by either kinase (Figure 4 C). This pointed to a second set of phospho-sites between amino acids 950-1144, consistent with SGD annotation (Figure 4 D).

**Figure 4.**
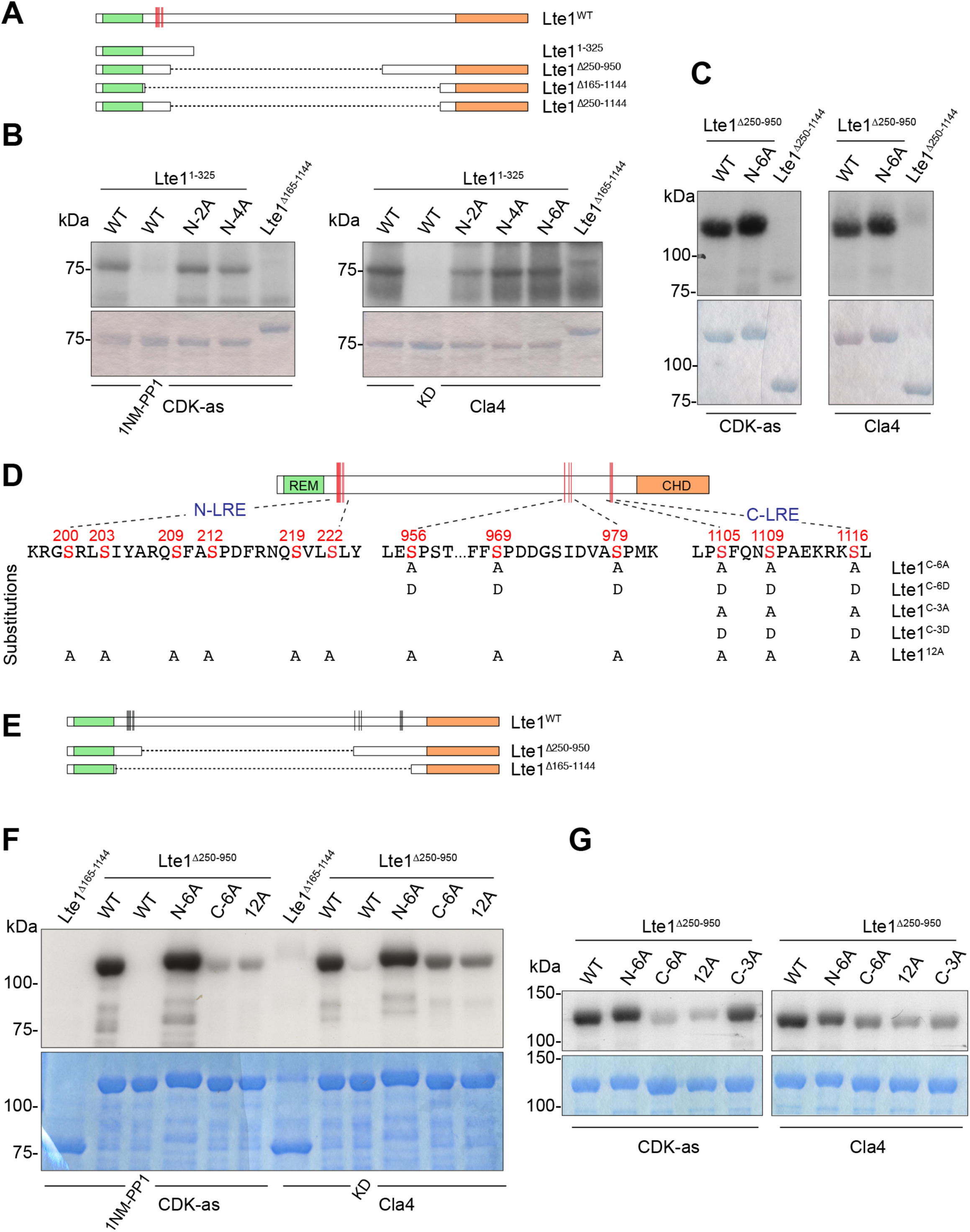
Failure of Cla4 and CDK to target the N-LRE *in vitro* pointed to further phosphorylation sites in a region near the CHD. (A-C) The N-LRE is not a prime target of Cla4 and CDK *in vitro*. (A) Deletion constructs used as kinase substrates. (B-C) *In vitro* kinase assay was carried out using the indicated substrates and either analogue-sensitive Clb2-Cdc28-as complex (CDK-as) or Cla4. After SDS-PAGE, the gel was stained with Coomassie blue and dried (lower panel) followed by exposure to film to detect radiolabeled species (upper panel). (B) Specific phosphorylation was detected on Lte1^1-325^ fragment but phosphorylation was unaffected by substitutions that cancelled the studied phosphorylation sites at the N-LRE. An internally deleted Lte1 substrate essentially consisting of the REM and CHD (Lte1^Δ165-1144^) showed reduced phosphorylation with either kinase. 1NM-PP1, inhibitory analog for CDK-as. KD, kinase dead version of Cla4. (C) Lte1^Δ250-950^ was phosphorylated by CDK-as or Cla4 irrespective of substitutions cancelling phosphorylation at the N-LRE. Yet, a further deletion eliminating sequences adjacent to the CHD in Lte1^Δ250-1144^ markedly reduced phosphorylation by both kinases. (D) Diagram outlining the position of regulatory sequences near the CHD. Phospho-sites annotated in the SGD and selected for genetic analysis are shown in red. Substitutions introduced in this study and their designation are indicated. The CHD-proximal cluster was referred to as the C-LRE here. (E-G) CDK and Cla4 target clusters of sites near the CHD (E) Deletion constructs used as kinase substrates. (F-G) *In vitro* kinase assays were performed with the indicated substrates and enzymes. After SDS-PAGE, the gel was stained with Coomassie blue and dried (lower panel) followed by exposure to film to detect radiolabeled species (upper panel). (F) In the context of the indicated substrates, 6 S to A substitutions significantly reduced phosphorylation by either CDK or Cla4 *in vitro*. Combining those with substitutions within the N-LRE did not affect phosphorylation further. 1NM-PP1, inhibitory analog for CDK-as. KD, kinase dead version of Cla4. (G) By contrast, only Cla4 phosphorylation was markedly reduced when S^1105^, S^1109^ and S^1116^ (C-3A; including one consensus PAK site) were substituted to A.

In the context of Lte1^Δ250-950^, S to A substitutions at S^956^, S^969^, S^979^, S^1105^, S^1109^, and S^1116^ (C-6A) reduced phosphorylation by either CDK or Cla4 *in vitro* (Figure 4 E and F) without further decrease when adding substitutions in the N-LRE (12A). Phosphorylation by Cla4 was similarly reduced when substitutions were introduced at S^1105^, S^1109^, and S^1116^ only (C-3A) but phosphorylation by CDK continued (Figure 4 G). Taken together, the sub-cluster closest to the CHD, referred to as C-LRE here was a prime target of Cla4 and contained one full PAK consensus site (R/K-R-X-pS/T). Further CDK sites were identified near this element.

### Phospho-mimetic substitutions at the C-LRE restored Lte1 recruitment to the bud cell cortex in a *cla4Δ* mutant

To explore the role of phosphorylation at the C-LRE we first expressed a mutant containing 6 phospho-mimetic substitutions (Lte1^C-6D^; Figure 4 D) fused to GFP in otherwise wild type cells. The protein was correctly confined to the bud cortex compartment (Suppl. Figure 3 A). Yet, the significance of part of these phospho-sites became apparent when Lte1^C-6D^-GFP and Lte1^C-3D^-GFP (Figure 4 D) were expressed in a *cla4Δ* background (Figure 5 A-C). By contrast to the limited Lte1^WT^-GFP cortical recruitment in *cla4Δ* cells, Lte1^C-6D^-GFP and Lte1^C-3D^-GFP marked the entire bud cortex (Figure 5 A and B). Single-cell linescan analysis confirmed that both Lte1^C-6D^ and Lte1^C-3D^ retained localization in a *cla4Δ* background, while bud cortical spreading scaled with bud growth throughout (Figure 5 C). Yet, maximal mean intensity in small budded cells was below that reached by Lte1^WT^ in otherwise wild type cells (compare with Figure 1 D).

**Figure 5.**
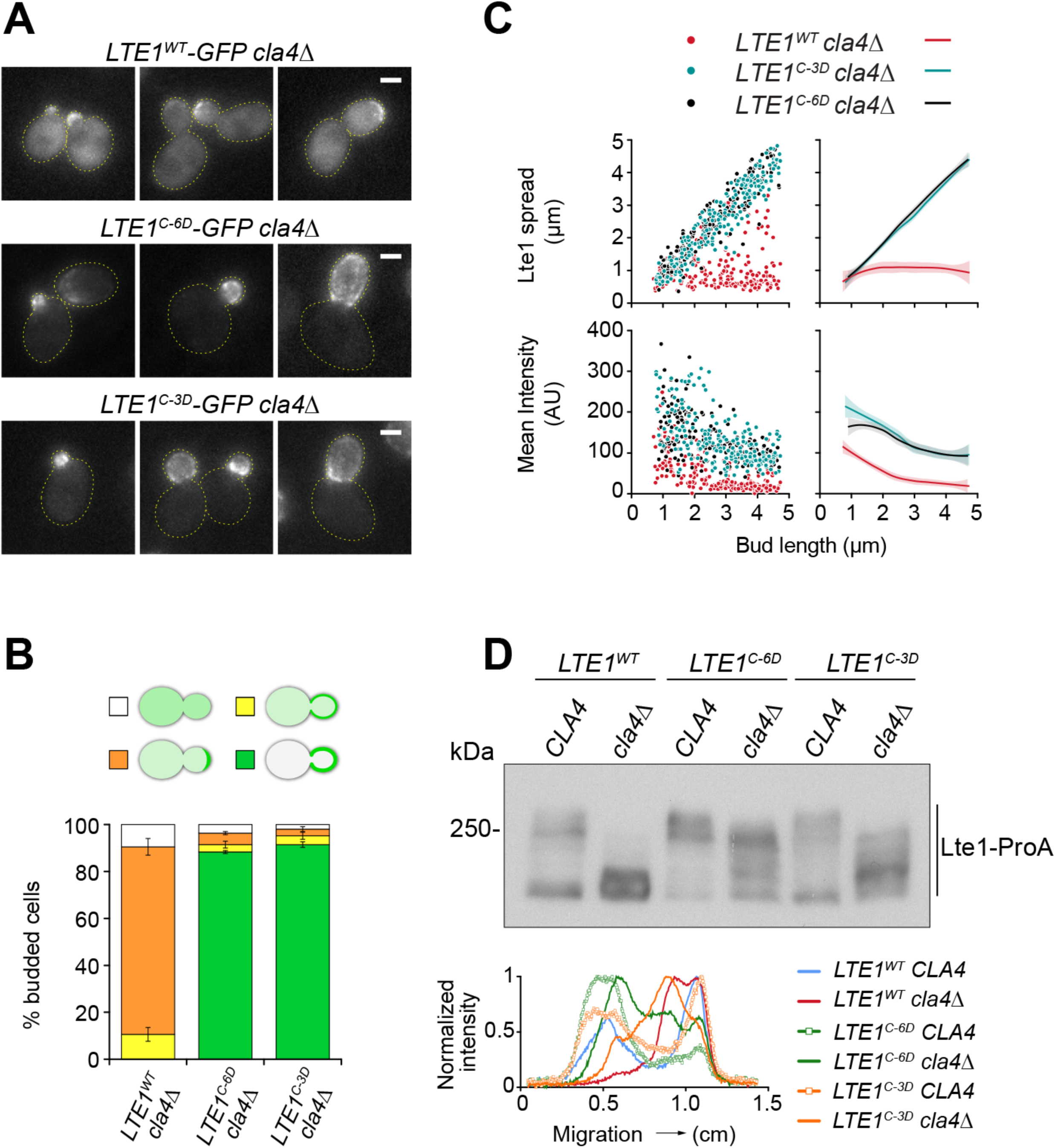
Phospho-mimetic substitutions at the C-LRE restored correct Lte1 cortical localization in a *cla4Δ* background. (A) Representative fluorescence images for localization of Lte1^WT^, Lte1^C-6D^ or Lte1^C-3D^ fused to GFP in a *cla4Δ* strain. Substitutions within the C-LRE rescued Lte1 localization in a *cla4Δ* background. Scale bar, 2 µm. (B-C) *LTE1^C-3D^* and *LTE1^C-6D^* suppressed the localization defect in a *cla4Δ* background to a comparable extent. (B) Distribution of modes of cortical localization in budded cells. The mean ± SEM of three independent counts of at least 120 cells is shown. Rescue by a single phospho-mimetic substitution at the PAK consensus site in the C-LRE is shown in Suppl. Figure 3 B-C. (C) Scatterplot (left) and smoothed plot (right) for Lte1 cortical spread and mean intensity in the bud as a function of bud length in the indicated strains (*LTE1^WT^ cla4Δ* n = 221, *LTE1^C-6D^ cla4Δ*, n =212, *LTE1^C-3D^ cla4Δ* n= 203). Solid lines correspond to the smoothed conditional mean and shading represents the 95% CI. The cumulative distribution of bud length in the populations analyzed is shown in Suppl. Figure 3 D. (D) Impact of phospho-mimetic substitutions on Lte1’s *in vivo* phosphorylation, assessed by western blot and linescan analyses. Whole extracts from the indicated strains were prepared after cell synchronization at metaphase. Phospho-mimetic substitutions at the C-LRE significantly restored Lte1 mobility shift in a *cla4Δ* mutant background. Yet, a contribution of Cla4-dependent mobility remains, suggesting the presence of additional sites outside the C-LRE or an indirect contribution to the profile.

Lte1^C-6D^ and Lte1^C-3D^ were hyper-phosphorylated *in vivo* contrary to Lte1^WT^ when expressed in a *cla4Δ* background, as indicated by manifest mobility-shift observed by western blot analysis (Figure 5 D). Yet, Cla4-dependent phosphorylation was still detected (compare the upper-most bands in *CLA4* vs *cla4Δ* lanes) either due to Cla4 sites outside the C-LRE or other indirect contribution by Cla4.

A phospho-mimetic substitution at the sole PAK consensus site within the C-LRE (Lte1^1116D^), also restored Lte1 cortical localization in a *cla4Δ* background (Suppl. Figure 3 B-C). Accordingly, average linescan traces of cells expressing Lte1^1116D^ closely resembled those of cells expressing Lte1^C-3D^ (Suppl. Figure 3 E). In both cases, the initial peaks corresponding to label in small budded cells were reduced relative to Lte1^WT^ in wild type cells (compare Suppl. Figure 3 E with Suppl. Figure 1 E). The incomplete rescue is consistent with previous proposals that Cla4 might help dock Lte1 at the cell cortex by physical interaction or by other Cla4-dependent anchors (Seshan et al., 2002).

In conclusion, phospho-mimetic substitutions within the C-LRE bypassed dependency on Cla4 (with a single substitution at a PAK site being sufficient for rescue) without otherwise disrupting compartmentalization of Lte1 to the bud. Furthermore, other protein kinases may phosphorylate Lte1 once recruited to the cell cortex and/or in response to priming phosphorylation by Cla4 at the C-LRE.

### A MARK/PAR-1-like kinase targets N-LRE sites

The fact that Lte1^C-6D^ regained bud cortex association together with a phospho-dependent mobility shift in *cla4Δ* cells suggested the participation of additional bud-localized protein kinases targeting the N-LRE.

Yeast Kin1 and Kin2 are members of the MARK/PAR-1 family of S/T protein kinases (Wu and Griffin, 2017) implicated in control of cell polarity. Kin2 is enriched at sites of polarized growth and interacts with the polarisome component Pea2, the septin Cdc11 and Rho3 (Yuan et al., 2016). Kin1 and Kin2 are functional homologues that share extensive sequence identity (Levin et al., 1987) and the consensus motif N-X-pS-X-S/T-X-I/L, which includes a proposed phospho-priming site at position +2 targeted by an unknown kinase (Jeschke et al., 2018). The N-LRE contains a close match to this consensus, pointing to the possibility that Kin1 and Kin2 might regulate Lte1 (Figure 6 A). Thus, localization studies were conducted after introducing phospho-mimetic substitutions at the putative Kin1/2 site in the N-LRE. The resulting Lte1^219D, 222D^-GFP fusion expressed in otherwise wild type cells was localized to the cortex of unbudded cells as well as the mother cell cortex of budded cells. Modes of localization resembled those observed for Lte1^N-4D^ but the bias to the mother cell was less pronounced (Figure 6 B).

**Figure 6.**
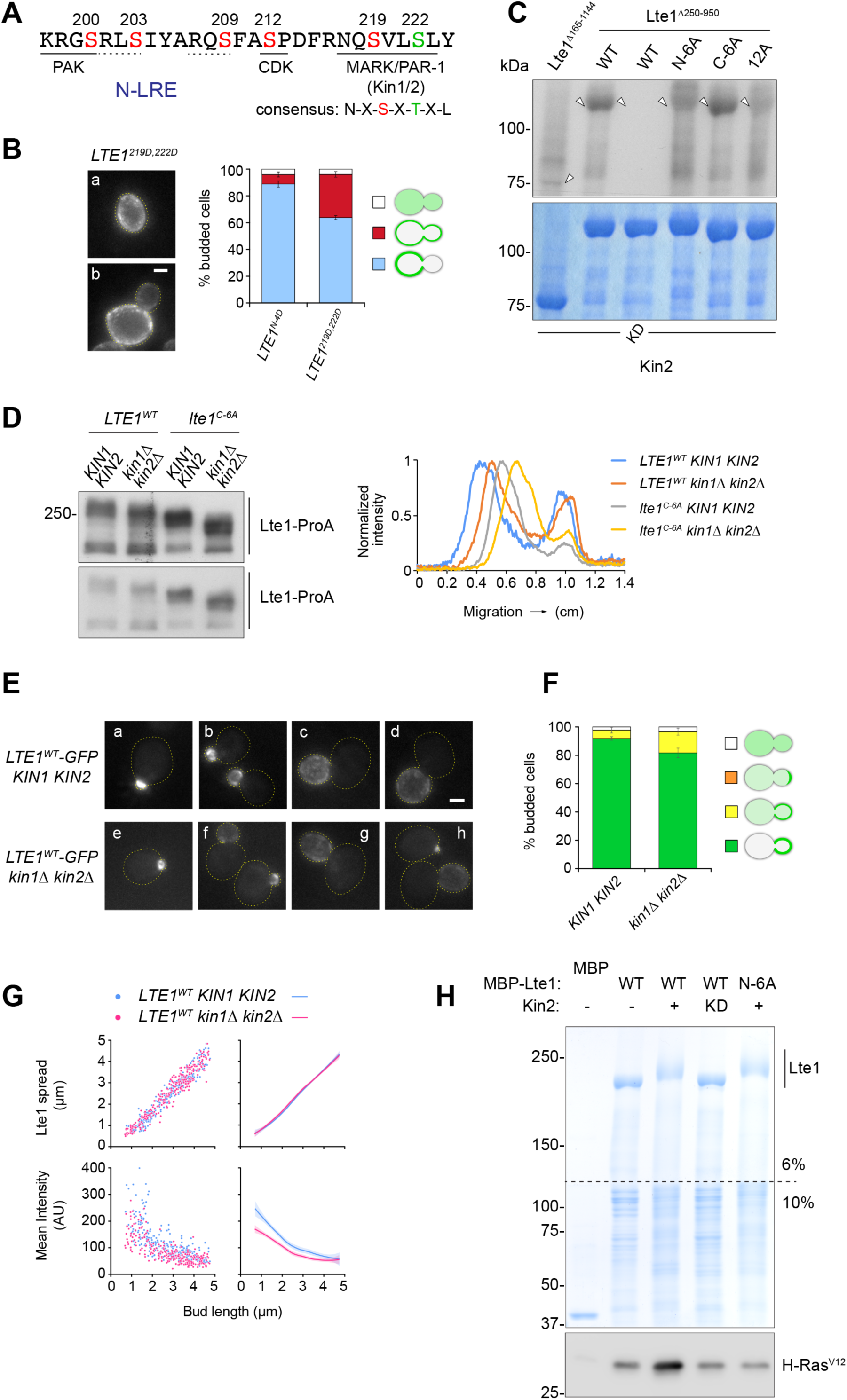
The MARK/PAR-1 like kinase Kin2 targets sites within the N-LRE. (A) N-LRE sequence outlining a potential site for MARK/PAR-1 like kinases. A PAK consensus site, 2 minimal Cla4 consensus sites (dotted line) and a minimal CDK consensus site are also shown. (B) Phospho-mimetic substitutions at a putative target site for Kin1/2 protein kinase within the N-LRE (S219D-S222D) directed Lte1 to the mother cell cortex in unbudded (a) or budded (b) cells (left panel). Modes of localization scored in budded cells for Lte1^N-4D^-GFP vs Lte1^219D,222D^-GFP (right panel). The plot shows mean ± SEM of three independent counts of at least 120 cells. Bias to the mother cell cortex was less pronounced for Lte1^219D,222D^ compared to Lte1^N-4D^. (C) *In vitro* kinase assay was carried out using the indicated substrates and either purified Kin2 or Kin2 kinase-dead mutant (KD). After SDS-PAGE, the gel was stained with Coomassie blue and dried (lower panel) followed by exposure to film to detect radiolabeled species (arrowheads, upper panel). Substitutions that cancelled phosphorylation within the N-LRE reduced phosphorylation by Kin2 while Lte1^C-6A^ was unaffected. (D) Impact of deletion of *KIN1* and *KIN2* on phospho-dependent mobility shift of Lte1-ProA vs Lte1^C-6A^-ProA assessed by western blot (left panel) and linescan analyses (right panel). Extracts were prepared from cells synchronized at metaphase. Two exposures of the blot are shown. (E) Representative images for localization of Lte1^WT-GFP^ in wild type (a-d) vs *kin1Δ kin2Δ* (e-h) budded cells. Scale bar, 2 µm. (F) Modes of Lte1^WT^-GFP localization collectively scored in budded cells in otherwise wild type vs *kin1Δ kin2Δ* cells. Data corresponds to mean ± SEM of three independent counts of at least 120 cells. (G) Scatterplot (left) and smoothed plot (right) for Lte1^WT^ cortical spread and mean intensity in the bud as a function of bud length in otherwise wild type or *kin1Δ kin2Δ* backgrounds (*KIN1 KIN2* n = 259, *kin1Δ kin2Δ* n = 246). Solid lines correspond to the smoothed conditional mean and shading represents the 95% CI. Overall, cortical recruitment was reduced without otherwise preventing cortical spreading. (H) Phosphorylation by Kin2 enhanced Lte1’s interaction with Ras-GTP *in vitro*, dependent on an intact N-LRE. MBP-Lte1^WT^ or MBP-Lte1^N-6A^ were substrates for either mock (-) or Kin2 phosphorylation (+) reactions before incubation with purified H-Ras^V12^-myc. A control reaction with a kinase-dead mutant version of Kin2 (KD) was included. After resolving pulled-down products by SDS-PAGE the lower part of the gel was excised and subjected to western blot analysis to visualize bound Ras while the top part was directly stained by Coomassie blue to view MBP-Lte1 fusion baits.

Supporting the idea of direct control, purified Kin2 but not Kin2^KD^ (kinase-dead) phosphorylated Lte1^Δ250-950^ *in vitro* (Figure 6 C). The phosphorylated band was reduced when substitutions cancelling phosphorylation sites in the N-LRE were introduced (Figure 6 C, N-6A or 12A) but was retained when such substitutions were introduced at the C-LRE only (Figure 6 C, C-6A). Taken together, Kin2 phosphorylates Lte1 at the N-LRE.

Then, the contribution of Kin1/2 to bulk Lte1 phosphorylation *in vivo* was assessed by western blot analysis of whole extracts from cells expressing Lte1^WT^-ProA or Lte1^C-6A^-ProA in wild type or *kin1Δ kin2Δ* background (Figure 6 D). A subtle decrease in mobility shift was observed for Lte1^WT^-ProA expressed in *kin1Δ kin2Δ* background. Lte1^C-6A^-ProA also exhibited a reduced mobility shift compared to Lte1^WT^-ProA which was further reduced when expressed in *kin1Δ kin2Δ* cells. Thus, Kin1/2-dependent phosphorylation and substitutions at the C-LRE were additive (Figure 6 D).

Conversely, Lte1^219D, 222D^-ProA retained a phospho-dependent mobility shift in a *cla4Δ* background, in line with the connection between restored cortical association and phosphorylated state independent of Cla4 (Suppl. Figure 4 A).

Lte1^WT^-GFP localization was mildly impaired in *kin1Δ kin2Δ* cells (Figure 6 E and F), an observation confirmed by single-cell linescan analysis (Figure 6 G). While the spread of the label continued to scale with bud growth, an overall reduction in mean label intensity at the bud cell cortex occurred. Average linescan traces also reflected this decrease, pointing to significant, yet redundant, contribution by Kin1/2 to the program (Suppl. Figure 4 C).

Since purified Kin2 could phosphorylate N-LRE sites and Lte1^219D, 222D^ was mislocalized to the mother cell cortex, the impact of Kin2 phosphorylation on Lte1 binding to Ras-GTP was tested. To this end, purified MPB-Lte1 was treated with Kin2 or Kin2^KD^ prior to the pull-down assay. Only phosphorylation of the bait by active kinase increased the amount of Ras-GTP recovered by pull-down (Figure 6 H).

Furthermore, MBP-Lte1^N-6A^ treated in the same way showed no increase in Ras-GTP signal (irrespective of additional phospho-sites apparent in full-length Lte1), suggesting that Kin2-dependent enhancement was mediated by phosphorylation at the N-LRE. Accordingly, Lte1^219D, 222D^ also showed enhanced binding to Ras-GTP *in vitro* (Suppl. Figure 4 D).

In conclusion, these data identified Kin1/2 as new protein kinases involved in the regulation of Lte1 binding to Ras-GTP via the N-LRE and its phospho-dependent program for cortical localization.

### Substitutions that cancel phosphorylation sites in Lte1 have a cumulative impact on Lte1 localization and function

Substitutions that prevented phosphorylation at the N- or C-LRE, had modest effect on Lte1^WT^ localization or function. By contrast, the impact of phospho-mimetic substitutions indicated that LREs support distinct mechanisms driving the program for Lte1 cortical localization. In an attempt to reconcile these observations, 12 substitutions that eliminated phosphorylation at the N-LRE, C-LRE and adjacent CDK target sites were combined (Figure 4 D). However, localization of Lte1^12A^-GFP in otherwise wild type cells was only mildly impaired (Figure 7 A and B). Similarly, S/T to A substitutions at all full-consensus CDK sites within Lte1’s central domain, which significantly reduced the cell cycle-dependent profile of Lte1 phosphorylation, failed to impair localization in a previous study (Jensen et al., 2002). Thus, a further mutant was generated by adding S/T to A substitutions at CDK sites in the central domain (T^317^, T^614^, S^630^, S^667^, T^793^ and S^854^). The resulting construct, Lte1^18A^-GFP, was initially localized at the incipient bud like Lte1^WT^ or Lte1^12A^. However, cortical label only persisted at the apical portion of the growing bud (Figure 7 A and B). Single-cell linescan analysis confirmed the deviation from the Lte1^WT^ program in cells expressing Lte1^18A^-GFP (Figure 7 C and Suppl. Figure 5 A-B). Recruitment to the incipient bud site was modestly affected and label retained in small budded cells. Yet, mean intensity further declined with bud growth, as the spread narrowed. Overall, it seemed that the bud cortical domain was not sufficiently established. Taken together, multisite phosphorylation over the budded interval of the cell cycle may govern Lte1 recruitment and retention to configure a bud cortical domain that should persist until the end of mitosis.

**Figure 7.**
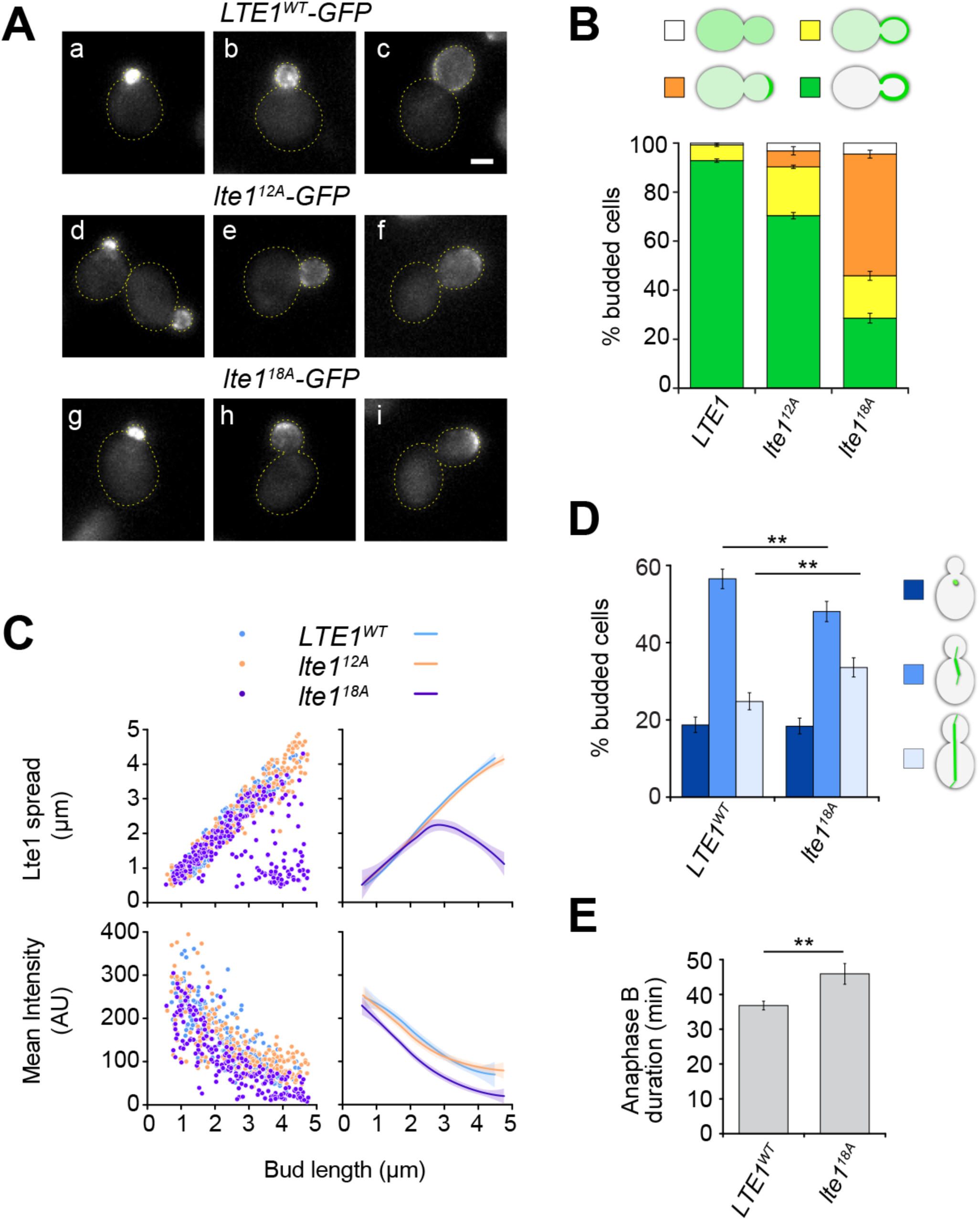
Cumulative effect of phospho-site substitutions on Lte1 cortical program. (A) Representative fluorescence images for localization of Lte1^WT^, Lte1^12A^ (see Figure 4 D) and Lte1^18A^ (additionally containing A substitutions at 6 previously identified CDK phosphorylation sites: T^317^, T^614^, S^630^, S^667^, T^793^ and S^854^; Jensen et al., 2002) fused to GFP, expressed in a wild type background. Scale bar, 2 µm. (B) Modes of Lte1 localization scored in budded cells for the same strains presented as mean ± SEM of three independent counts of at least 120 cells. (C) Scatterplot (left) and smoothed plot (right) for Lte1^WT^-GFP, Lte1^12A^-GFP and Lte1^18A^-GFP cortical spread and mean intensity in the bud as a function of bud length in otherwise wild type cells (*LTE1^WT^* n = 272, *lte1^12A^* n = 331, *lte1^18A^* n= 283). Solid lines correspond to the smoothed conditional mean and shading represents the 95% CI. The cumulative distribution of bud lengths in each population is presented in Suppl. Figure 5 A. Lte1^18A^ was impaired for overall recruitment and progressively retained limited apical localization with bud growth. (D-E) *lte1^18A^*cells exhibited a mitotic exit delay. (D) Distribution of budded cells along the spindle pathway assessed in still images from asynchronous populations of *LTE1* (n = 1509 cells) or *lte1^18A^* (n = 1427 cells) strains. The *lte1^18A^*strain showed a 27% increase in elongated-to-short spindle ratio, consistent with the lengthening of the anaphase interval. Error bars show 95% CI. ** p<0.01 according to the two proportion Z-test. (E) Mean duration of anaphase B was determined in *LTE1* (n = 30) vs *lte1^18A^* (n = 21) cells by time lapse analysis as described in Materials and Methods. Error bars show SEM. ** p<0.01 according to the Mann Whitney two-tailed test.

The fact that Lte1^18A^ localized apically in large budded cells suggests that perhaps the boundary between inhibitory and permissive domains framing the SPOC in late mitosis might be shifted, thus delaying sensor activation upon SPB entry in the bud due to the reduction in the domain marked by Lte1. To investigate this possibility, mitotic spindle progression was analyzed in *LTE1^WT^* or *lte1^18A^* cells expressing GFP-Tub1 (tagging α-tubulin). First, the distribution of budded cells by mitotic spindle stage was assessed in still images of asynchronous cultures (Figure 7 D). Relative to *LTE1^WT^*, *lte1^18A^* increased cells with elongated spindles, indicating a longer anaphase interval. Indeed, time lapse analysis showed that anaphase duration was increased by 25 % in *lte1^18A^*cells (Figure 7 E), confirming that Lte1^18A^ failed to promote timely mitotic exit. Still, *lte1^18A^* rescued *lte1Δ spo12Δ* co-lethality indicating a partial loss-of-function (Suppl. Figure 5 C).

Taken together, a combination of substitutions preventing phosphorylation of multiple regulatory sites in Lte1 significantly disrupted the program for demarcation of the bud compartment. The fact that *lte1^18A^*continued to rescue known *lte1Δ* phenotypes (co-lethality with *spo12Δ*) emphasizes the significance of correctly patterning Lte1 cortical association *per se* for timely mitotic exit in cycling cells.

### Differential effect of phospho-mimetic substitutions at N-LRE vs C-LRE on Lte1-Ras2 complex formation *in vivo*

To complement the analysis for the impact of substitutions within regulatory sites on the interaction between Lte1 and Ras-GTP, the ability of Lte1 to pull down Ras-GTP *in vitro* was determined using MBP-Lte1^WT^ vs MBP-Lte1^18A^, MBP-Lte1^N-6A^ or MBP-Lte1^C-6D^ as baits (Figure 8 A). We found that the inherent ability of Lte1 to bind Ras-GTP *in vitro* remained intact. Furthermore, MBP-Lte1^C-6D^ did not show enhanced binding to Ras-GTP, contrasting with the effect of phospho-mimetic substitutions within the N-LRE (Figure 3).

**Figure 8.**
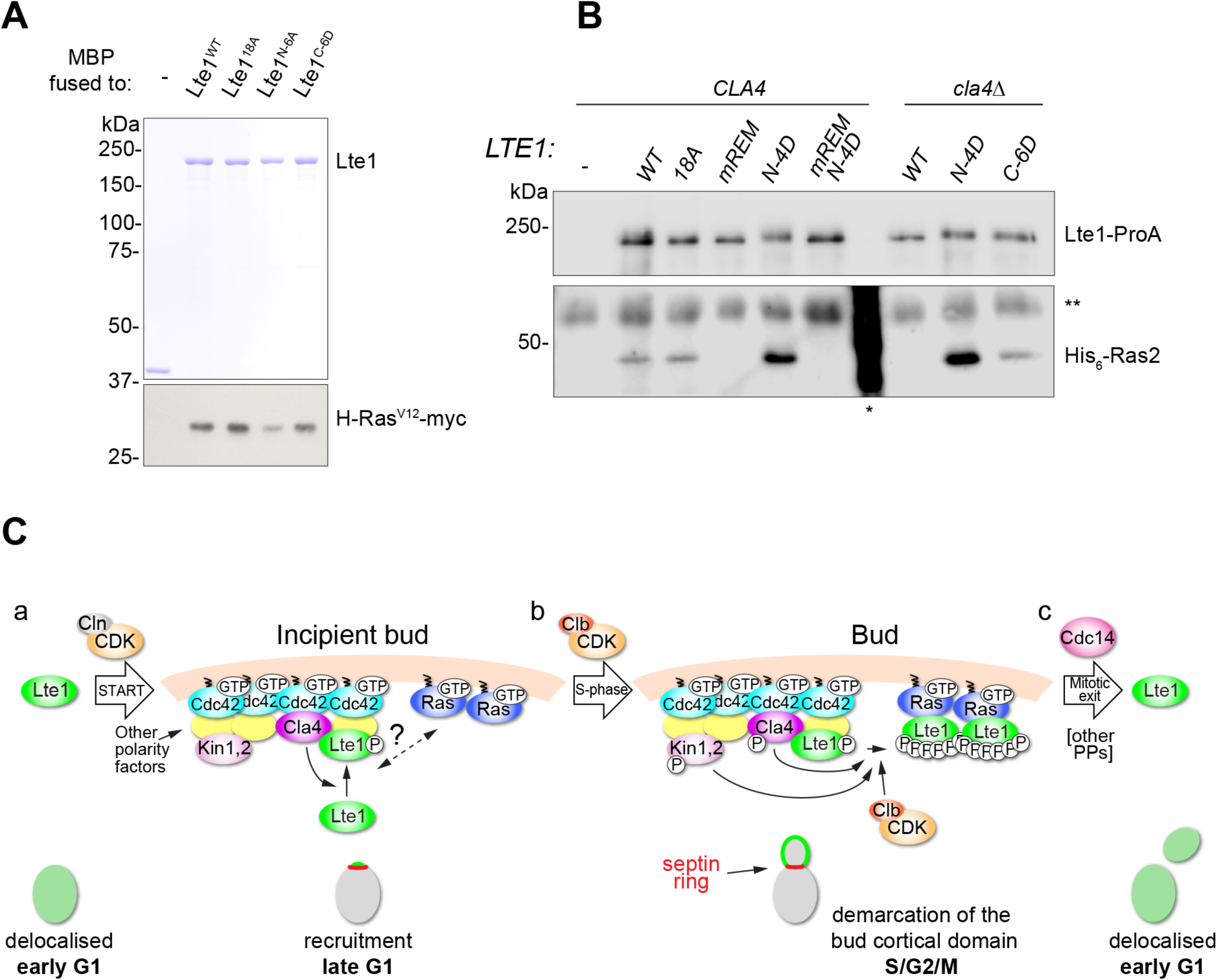
Control of Lte1-Ras complex formation *in vivo* by N-LRE vs C-LRE. (A) *In vitro* pull-downs performed as described in Figure 3 F. After resolving the pull-down product by SDS-PAGE the lower part of the gel was excised and subjected to western blot analysis to visualize bound Ras while the top part was directly stained by Coomassie blue to view control MBP or MBP-Lte1 fusion baits. Lte1’s inherent ability to pull down Ras-GTP *in vitro* was unaffected by combining substitutions that cancel phosphorylation at N-LRE, C-LRE and CDK sites studied here. In addition, phospho-mimetic substitutions in Lte1^C-6D^ did not enhance binding to Ras-GTP in contrast to the effect such substitutions had at the N-LRE (Figure 3 F). (B) *In vivo* interaction between Lte1 and Ras2 assessed by coimmunoprecipitation. Whole extracts were prepared from metaphase-arrested cells expressing the indicated versions of Lte1 fused to ProA and His_6_-Ras2 replacing the respective endogenous loci in wild type or *cla4Δ* backgrounds. Co-precipitated Lte1 and Ras2 were analyzed by western blot. [*] MW standard reacted with anti-His antibody; [**] IgG heavy chain cross-reacting band. (C) A model for multisite phosphorylation confining Lte1 to the bud cortical compartment. (a) In early G1, Lte1 is unphosphorylated and delocalized. G1-CDK (Cln-CDK) triggers passage through START. In turn, local activation of Cdc42 at the site selected for bud emergence dictates the recruitment of various polarity determinants including the PAK Cla4. Polarity factors and Cla4 may redundantly tether Lte1 while it undergoes phosphorylation at the C-LRE. These events would prime Lte1 recruitment to the incipient bud cell cortex. Kin1,2 enrichment at the emerging bud may also involve polarity factors such as Pea2 and Kel1 (Breitkreutz et al., 2010; Yuan et al., 2016). Whether Ras contributes at this stage of recruitment is unclear (?). (b) Following entry into S phase, Clb-CDK targets Kin1,2 and Cla4 (Holt et al., 2009; Lanz et al., 2021), thus contributing indirectly as well as directly to hyper-phosphorylation of primed Lte1 at and away from the N-LRE. This translates into Lte1 persistent association with Ras-GTP exclusively within the bud throughout the rest of the cell cycle. (c) Upon mitotic exit Lte1 dephosphorylation by Cdc14 and PP2 phosphatases (Touati et al., 2019), reverses both binding to Ras and cortical association.

We then pursued the validation of the role of phospho-regulatory sites in formation of an Lte1-Ras complex *in vivo* assessed by immunoprecipitation using whole extracts from cells expressing epitope-tagged fusions of wild type or mutant Lte1 and yeast Ras2 at endogenous levels (Figure 8 B).

Ras2 was co-immunoprecipitated with Lte1^WT^-ProA or Lte1^18A^-ProA but was not recovered upon immunoprecipitation of Lte1^mREM^-ProA. This result confirmed the critical role of the REM in supporting cortical localization dependent on Ras2 binding *in vivo.* However, Ras2 co-immunoprecipitation with Lte1^18A^ seemed unaffected, perhaps due to poor signal dynamic range to discriminate a reduction from wild type binding level. Yet, supporting the link between constitutive cortical localization and enhanced binding to Ras, Lte1^N-4D^-ProA yielded markedly increased co-immunoprecipitated Ras2, dependent on an intact REM (compare WT, N-4D and N-4D-mREM).

A *cla4Δ* background prevented Ras2 co-immunoprecipitation with Lte1^WT^-ProA, as shown previously (Seshan and Amon, 2005). Interestingly, co-immunoprecipitation was restored differently in response to phospho-mimetic substitutions at the N-LRE vs the C-LRE. Indeed, Lte1^N-4D^-ProA co-immunoprecipitated Ras2 as efficiently as it did in a *CLA4* background. By contrast, Ras2 co-immunoprecipitation with Lte1^C-6D^-ProA matched the level observed for Lte1^WT^ in *CLA4* background. These data indicate that Lte1^N-4D^-Ras2 complex formation may be constitutive and refractive to Cla4 loss *in vivo*, due to sufficiently increased inherent affinity of Lte1 toward Ras-GTP from the outset (Figure 3 F and Suppl. Figure 2 E-F). By contrast, Lte1^C-6D^-ProA bypassed Cla4 but not further phospho-regulation to anchor Lte1 via Ras in the bud.

These results confirmed distinct mechanistic contributions for C vs N phospho-regulatory elements in initiating and establishing Lte1-Ras complexes, a critical sequence for enforcing exclusive Lte1 association with the bud cortical compartment.

## DISCUSSION

### Is the REM functionally conserved between Lte1 and hSOS1?

Lte1 cortical association via Ras requires sequences related to the REM and CHD of Ras-GEFs. Yet, only Ras-GTP can serve as a cortical anchor (Seshan and Amon, 2005; Yoshida et al., 2003). This selectivity implicated the Lte1 REM in Ras binding, by extrapolating from the role of this motif in hSOS1 (Margarit et al., 2003). To probe functional conservation, we characterized Lte1^mREM^, a mutant encoding substitutions analogous to those disrupting Ras-GTP binding to the hSOS1 REM (Sondermann et al., 2004).

Lte1^mREM^ failed to interact with Ras *in vitro* and *in vivo* (Figure 3 F and Figure 8 B). Accordingly, Lte1^mREM^ cortical localization and function were severely impaired and bulk phosphorylation reduced (Figure 2 E-G). Thus, the Lte1 REM is a critical determinant for cortical anchoring via Ras-GTP. Yet, Lte1 efficient interaction with Ras2 also involves the CHD (Geymonat et al., 2009; Yoshida et al., 2003).

The fact that the REM and CHD map to opposite ends of Lte1 and that the protein displays self-association between N- and C-terminal sequences (Jensen et al., 2002), could prove mechanistically relevant to Lte1 phospho-dependent anchoring via Ras-GTP, perhaps extending similarities between Lte1 and SOS. Indeed, activation of hSOS1 entails both recruitment to the cytoplasmic membrane through phospho-dependent docking to an activated receptor via adaptor proteins, and conformational change prompted by membrane association, which exposes the allosteric site for Ras-GTP binding. This would further tether hSOS1 to the membrane while stimulating GDP-GTP exchange by the CHD (Bandaru et al., 2019). In Lte1, these steps could be controlled by its own phospho-regulatory sites. Furthermore, the conserved role of the REM suggested here, may give support to a proposal by Seshan and Amon (2005) that binding of Ras-GTP to the REM might link Lte1 cortical localization with allosteric activation of Lte1-GEF, perhaps explaining the failure to detect Lte1-GEF activity towards Tem1 *in vitro* (Geymonat et al., 2009). Collectively, these appealing ideas remain to be explored.

### Polarized recruitment and anchoring via Ras are controlled by distinct Lte1 phospho-sites

Phosphorylation by Cla4 has been primarily implicated in Lte1 cortical association via Ras (Hofken and Schiebel, 2002; Seshan and Amon, 2005). Yet residual polarized localization of Lte1 in *cla4Δ* or even *ras1Δ ras2Δ* cells (Figure 1 and Suppl. Figure 2), suggests other intervening steps toward Lte1 cortical anchoring. Regarding CDK, cortical association and Ras binding may be uncoupled, according to cell cycle or developmental context. In cycling cells, G1-CDK is essential for Lte1 recruitment together with Cdc42 (Jensen et al., 2002). Overexpression of the CDK inhibitor Sic1 prevents Lte1-Ras coimmunoprecipitation despite Lte1 polarized cortical localization and partial phosphorylation. Lte1-Ras immunocomplexes are only detected after mitotic CDK activation (Geymonat et al., 2010; Seshan and Amon, 2005). Moreover, Lte1 concentrates at the shmoo tip independent of CDK during mating pheromone arrest, also unable to coimmunoprecipitate Ras (Geymonat et al., 2010).

To gain mechanistic insight into phospho-regulation and its links to Lte1-Ras complex formation, we first identified Lte1^Δ250-950^, a minimal deletion construct that preserved Lte1 program of localization (Figure 2 A-C). Lte1^Δ250-950^ retained two sequences we now refer to as N-LRE and C-LRE, with a number of potential phosphoregulatory sites that became the focus of our study. On the premise that their phosphorylation would be linked to Lte1 cortical association, we started by introducing phospho-mimetic substitutions in the context of full-length Lte1. We then pursued *in vitro* kinase assays to assign those key sites as targets for Cla4 or CDK. This analysis revealed contrasting roles for those two LREs and led to the identification of a new protein kinase participating in the program.

The C-LRE was a direct target for Cla4 phosphorylation (Figure 4). Phospho-mimetic substitutions in this element restored all features of the program in a *cla4Δ* mutant background. The substitution at the PAK site was sufficient for rescue (Suppl. Figure 3 B-E). Yet, phospho-mimetic substitutions did not enhance Lte1 binding to Ras *in vitro* nor *in vivo*, explaining why Lte1^C-6D^ remained responsive to polarizing cues (Figures 5 and 8). By contrast, phospho-mimetic substitutions within the N-LRE enhanced Lte1 binding to Ras, thus eliminating any spatio-temporal constraints for Lte1 association with the cell cortex, irrespective of Cla4 (Figures 3 F and 8 B). Interestingly, the extent of bias toward the mother cell cortex and ensuing SPOC defect were correlated with the strength of mutant Lte1 binding to Ras *in vitro*.

The N-LRE contains a minimal CDK consensus site and three potential Cla4 sites (R-X-pS/T, Mok et al., 2010), yet, we failed to detect their phosphorylation by either CDK or Cla4 *in vitro* (Figure 4). Perhaps our assay failed to reconstitute phosphorylation events that might otherwise occur *in vivo*. However, the fact that Lte1^C-6D^ expressed in a *cla4Δ* background retained compartmentalized localization to the bud and hyper-phosphorylation, suggested that additional protein kinases that might target the N-LRE to modulate binding to Ras might be enriched at sites of polarized growth. This led us to evaluate Kin2, one of a pair of redundant yeast MARK/PAR-1 kinases. Kin2 phosphorylation at N-LRE sites enhanced the interaction between Lte1 and Ras-GTP *in vitro* (Figure 6 H). Accordingly, phospho-mimetic substitutions at the presumed Kin1/2 site in the N-LRE also enhanced binding to Ras-GTP *in vitro* and drew Lte1 to the mother cell cortex (Figure 6 B and Suppl. Figure 4 D). Taken together, our studies identify Kin1/2 as one of the protein kinases that directly modulates Lte1 binding to Ras via the N-LRE.

The combination of substitutions precluding phosphorylation at N-LRE, C-LRE and six additional CDK sites along the central domain of Lte1 (Jensen et al., 2002) disrupted its program of localization. Lte1^18A^ showed limited apical localization in large budded cells. The fact that the *lte1^18A^* allele increased the duration of anaphase, suggested that the perturbed localization might impair Lte1 function as a timely activator of mitotic exit.

In conclusion, the use of phospho-mimetic substitutions has revealed distinct contributions for the C-LRE and N-LRE in Cla4-directed recruitment and Lte1-Ras complex formation, respectively. However, genetic analysis based on inactivation of phospho-sites pointed to redundancy in the program perhaps involving CDK and Cla4 alternative targets. Such multi-layered control and the apparent redundancy built into the program may explain the deployment of alternative cortical anchors for Lte1 by developmental or cell cycle context as stated above. This flexibility may extend to Lte1 role in MEN-independent control of cell polarity via the small GTPase Rsr1 (Geymonat et al., 2010).

### Lte1 integrates spatial and temporal control using multisite phosphorylation

Multisite phosphorylation and cross-compartment signal propagation are recurrent themes in cell cycle control (Almawi et al., 2020; Jones et al., 2018; Miller and Turk, 2018; Ord et al., 2019; Rock et al., 2013; Zhou et al., 2021). Phospho-sites may govern substrate interaction with a scaffold protein and/or recruitment to a subcellular localization, thus compartmentalizing further activation.

The inextricable relationship between Lte1 cortical association and phosphorylation has been attributed to interdependence between Cla4 and Ras (Hofken and Schiebel, 2002; Seshan and Amon, 2005). In that view, local activation of Cdc42 and Cla4 singles out the incipient bud for Lte1 phosphorylation and binding to Ras (against a backdrop of Ras over the entire cell membrane). Yet, our findings suggest a more convoluted set of combinatorial controls constraining Lte1 to the bud.

In this model (Figure 8 C), Lte1 functions as an integrator of multiple inputs, with each protein kinase contributing temporal and/or spatial cues. G1-CDK is the cell cycle trigger (Jensen et al., 2002), Cla4 may provide a cortical tether (Seshan and Amon, 2005) and create a primed substrate as pre-requisites for Lte1 accessing locally active protein kinases at sites of polarized growth like Kin1/2, perhaps acting in collaboration with mitotic-CDK and even mitotically active Cla4 (Benton et al., 1997). Conversely, Lte1 conformational changes linked to membrane binding might permit protein kinase access to defined target sites, thus accounting for interdependence between cortical association and phosphorylated status. The septin ring adds a diffusion barrier between compartments (Castillon et al., 2003; Hofken and Schiebel, 2002; Jensen et al., 2002; Seshan et al., 2002). These steps would allow and then reinforce Lte1-Ras complex formation only after a bud has formed, defining a domain for timely activation of mitotic exit. Thus, Lte1 is uniquely placed at the interface between cell cycle (Bardin et al., 2000; Pereira et al., 2000) and cell polarity (Geymonat et al., 2010), to impose coordination between the two at the M/G1 transition.

An outstanding question is how this timeline of phosphorylation may be interlaced with control of protein-protein interactions that, transiently or in concert with Ras, support Lte1 polarized recruitment. In this regard, the small GTPase Rsr1 and the polarity determinant Kel1 are significant interactors, even though neither is absolutely essential for Lte1 localization (Geymonat et al., 2010; Hofken and Schiebel, 2002; Jensen et al., 2002; Seshan and Amon, 2005). It will be possible to establish their precise contributions with the tools generated in this study.

Our studies have uncovered a novel type of phospho-dependent partnership by which Lte1 is positioned at the interface linking distinct small GTPase modules with cell cycle control, given its dual potential as effector (of Cdc42 and Ras) and regulator (of Rsr1 and Tem1). Many of these components are conserved in networks that control cell proliferation or cell polarity including PAKs, CDKs, MARK/PAR-1, Ras, Cdc42, and the Hippo pathway (Chetty et al., 2022; Phillips et al., 2024; Sells and Chernoff, 1997; Wu and Griffin, 2017). For example, Cdc42-mediated polarity and the Hippo tumor-suppressor pathway (showing conservation with the MEN kinase core module) may be part of a functional axis for correct balance between the intestinal stem cell and the transit-amplifying pools that supports tissue homeostasis (Zhang et al., 2022). Also, cross-talk between Ras signaling and the Hippo pathway has significant implications in tumorigenesis (O’Neill, 2017). Given the recognized relevance of these multiple players to proliferative disorders (Jang et al., 2020; Rane and Minden, 2019), achieving an integrative view, informed by studies in yeast, constitutes an important goal for uncovering new targets for therapeutic intervention.

## MATERIALS AND METHODS

### Yeast strains, constructs and genetic procedures

Yeast strains used in this study are listed in Supplementary Table 1 and were isogenic derivatives from 15DaubA *MAT***a** *bar1 ade1 his2 leu2-3,112 trp1-1a ura3Dns arg4* (Geymonat et al., 2020; Jensen et al., 2002; Juanes et al., 2013; Richardson et al., 1989), except when indicated otherwise. Yeast genetic procedures were according to Guthrie and Fink (1991). YEP or synthetic medium with supplements, contained 2% w/v dextrose or 1% w/v sucrose according to the experiment. Induction of P*_GAL1_*-driven expression was carried out in 1% w/v galactose as sole carbon source. Counter-selection of *URA3*-carrying plasmids was carried out in synthetic complete medium with 0.1% w/v 5-Fluoroorotic acid (5-FOA). Deletion of *LTE1, KIN1* and *KIN2* was carried out by one-step disruption using a targeted *KANMX, Hph or URA3* cassettes produced by PCR (Geymonat et al., 2010). Deletion of *RAS1* and *RAS2* genes was carried out by one-step disruption using the insert from plasmid pRA545 (Tatchell et al., 1984) and a targeted *TRP1* cassette produced by PCR, respectively. One-step disruption of *BCY1* was carried out using a targeted *ADE1* cassette. pHIS2-P_PGK1_-H-Ras was derived from pYGA-Ras (Clark et al., 1985). Strains expressing a GFP-Tub1 (yeast α-tubulin) fusion were created by transformation with pOB1 linearized with *Bsu*36I for integration at *TRP1* (Ten Hoopen et al., 2012).

*LTE1^WT^* or mutated versions encoding substitutions that either cancelled (S/T to A) or mimicked (S/T to D) phosphorylation at sites selected for study, were generated by nested PCRs. The resulting ORFs were inserted as 4.3 kb *Sal*I-*Not*I fragments to generate in-frame fusions to GFP expressed under the control of P*_HIS3_* in YIplac128. This plasmid series was used for transformation of a *lte1Δ* strain after linearization with *Bst*EII for integration at *LEU2*. Expression levels monitored by western blot analysis were found to be comparable for all mutants (Suppl. Figure 1 B). A modification of the intergenic flip flop one-step method (Mallet and Jacquet, 1996) was used to generate strains expressing *LTE1^WT^* or mutated versions fused to a protein A (ProA) tag under the control of the endogenous *LTE1* promoter. All constructs were verified by DNA sequencing.

### Digital imaging microscopy

Early-log cultures grown at 23 °C in synthetic complete medium supplemented with 0.1 g/L adenine were concentrated by centrifugation at 380 x g for 5 min and directly spotted onto slides for still-image acquisition of random fields of cells with a Nikon Eclipse E800 with a CFI Plan Apochromat 100x, NA 1.4 objective, a Chroma Technology GFP filter set and a Coolsnap-HQ CCD camera (Roper Scientific), controlled by MetaMorph software (Molecular Devices). Single DIC images at the middle focal plane were paired to 5-plane Z-stacks of fluorescence images obtained at 0.8 μm intervals between planes with 2 x 2 binning (Guo and Segal, 2017). Representative images were selected from maximal intensity projections obtained with the open-source program FIJI (Schindelin et al., 2012). After adjusting consistently brightness and contrast, and scaling to 8-bit image depth, cell images were incorporated into final figures using Adobe Illustrator.

Time lapse recordings were performed by mounting cells from logarithmic cultures in complete synthetic dextrose containing 25 % w/v gelatin. 5-Z fluorescence image stacks paired to a transmitted light image at the middle focal plane were acquired at 1- or 2-min intervals using a modified Nikon Eclipse E800 fitted with OptoLED heads (Cairns Research) and Chroma Technology filter sets, a CFI Plan Apochromat 100x, NA 1.4 objective, a Z-focus drive and an Orca-Flash 4.0 v2 CMOS camera (Hamamatsu), all controlled by MetaMorph software (Guo and Segal, 2017).

Measurements were performed using FIJI plugins on maximal intensity projections after subtracting image background. Linescans for Lte1-GFP fluorescence intensity along the cell polarity (mother-bud) axis were generated with the line tool set to 5-pixel width. Gaussian fit of the resulting profile peaks was used to derive one-dimensional mean integrated intensity and peak width at half maximal intensity in individual budded cells paired to the length of the bud axis independently measured in DIC images with the line tool. Values were imported into Microsoft Excel for plotting. Smoothed conditional mean plots with 95% CI were generated using the open-source package R and the function geom_smooth (method = loess). Additionally, linescan datasets were ordered by measured bud length and grouped at 0.5 µm intervals between 1 ± 0.25 µm and 4.5 ± 0.25 µm. Traces in each group were realigned with respect to the position of the bud neck for averaging. Modes of Lte1-GFP cortical label were scored in still images of random fields of cells according to the categories outlined in Suppl. Figures 1 and 2. Data corresponded to the mean of three independent counts of at least 120 cells. Quantification of Lte1-GFP mean label intensity in time lapse series was carried out on maximal intensity projections by generating a binary mask in order to segment cortical label over the entire bud. The resulting regions of interest were reviewed and, if necessary, edited manually. Measurements were collated for plotting in Microsoft Excel.

The mitotic spindle was visualized by still-imaging or time lapse in cells expressing GFP-Tub1. The distribution of budded cells according to mitotic spindle stage was scored in maximal intensity projections of still-images for random fields of asynchronous cells obtained as outlined above. Spindle stages in budded cells were categorized as follows: a) unseparated spindle poles (single focus of label associated with cytoplasmic microtubules); b) short spindles (from spindle pole separation to 2.5-µm long spindle); c) elongated spindles (> 2.5 µm long to fully elongated). The duration of anaphase B was determined in time lapse series as the interval from onset of the fast phase of spindle elongation until the last plane in which the elongated spindle was intact. Cells that did not reach spindle disassembly within the recorded time were not included.

### Lte1_REM-CHD_ model

Lte1 (residues 1-185 and 1190-1435) with two molecules of Ras1 (residues 8-174) were modelled based on the Ras-Sos1 complex (PDB 1nvv) with *Modeller* V9.20 (Sali and Blundell, 1993) and using slow loop refinement.

### Protein production and purification

MBP fusions to full-length or Lte1 fragments of wild type or phospho-site mutant versions were expressed and purified from yeast using an auto-selection expression system (Geymonat et al., 2007). Briefly, PCR cassettes encoding the relevant ORF flanked by homologous sites to the ends of gapped pMG3 expression vector were used for transformation of the host MGY853 (*MAT a ura3-1 trp1-28 leu2*Δ*0 lys2 his7 cdc28::LEU2 pep4::LEU2 [URA3-CDC28])* by gap-repair. Transformants were plated on 5-FOA to counter-select the resident *CDC28*-carrying plasmid. From that point, the expression construct was maintained in rich medium allowing for efficient protein expression. The strains used for expression of MBP-Lte1 fusion constructs are included in Supplementary table 1.

Protein induction was carried out in YEP-galactose for 8 hours at 30 °C. Cells were harvested, rinsed with ice-cold water, snap-frozen and stored at -80 °C. Pellets were resuspended to 1/20 volume of the original cell culture in cold breaking buffer (50 mM Tris-HCl pH 7.5, 250 mM NaCl, 0.2% NP-40, 10% v/v glycerol, 5 mM EDTA, Roche cOmplete-EDTA free protease inhibitor, 2mM PMSF, 5 mM DTT) and disrupted in a FastPrep-24 grinder (MP Biomedicals) using 0.5-mm glass beads. Cell lysates were cleared by two cycles of centrifugation at 16,000 x g for 5 min at 4 °C. Amylose resin (New England Biolabs) was added and allowed to bind for 2 h on a rotator at 4 °C. The bound amylose resin was washed 6 times with washing buffer (50 mM Tris-HCl pH 7.5, 250 mM NaCl, 0.2% NP-40, 1 mM DTT) and once with 50% v/v glycerol in PBS. A ∼500 µL slurry of bound resin in 50% v/v glycerol in PBS was stored at -20 °C. A 20 µL aliquot was resolved by SDS-PAGE and bound protein quantified by Coomassie blue staining against a BSA standard curve.

Purification of constitutively active Ras-GTP used in pull-down assays *in vitro* was undertaken using plasmid pMH919 encoding His_6_-H-Ras^V12^-myc (strain MGY905, Supplementary table 1). As expression of Ras2^V19^ was incompatible with the *GAL* induction protocol applied here, we took advantage of the fact that H*-ras* can fully replace yeast *RAS* genes for Lte1 localization (Suppl. Figure 2). The purification of the His-tagged protein was carried out following the above steps but using a different breaking buffer (50 mM Tris-HCl pH 7.5, 250 mM NaCl, 0.2% NP-40, 10% v/v glycerol, 20 mM imidazole, 5 mM β-ME, Roche cOmplete-EDTA free protease inhibitor, 2 mM PMSF), nickel resin (Super Ni-NTA Affinity Resin, Generon, #NB-45-00042-25) and washing buffer (50 mM Tris-HCl pH 7.5, 250 mM NaCl, 0.2% NP-40, 40 mM imidazole), followed by addition of elution buffer (30 mM Tris-HCl pH 7.5, 200 mM NaCl, 200 mM imidazole) to the packed resin to release soluble Ras protein. Protein concentrations were measured with Bradford reagent, and the most concentrated aliquots were combined for overnight dialysis against buffer A (20 mM Tris-HCl pH 7.5, 150 mM NaCl, 10% v/v glycerol, 500 µM DTT). Dialyzed proteins were assessed by SDS-PAGE and final aliquots stored at -80 °C.

A Clb2^Δdb^ stabilized version of the mitotic cyclin Clb2 was produced as a FLAG-Clb2^Δdb^-Chitin-Binding Domain (CBD) fusion protein in the context of overexpressed analog-sensitive Cdc28-as/His_6_-Cks1 (strain MGY1111, Supplementary table 1). The CBD/intein tag enabled one-step purification by binding to chitin-resin (New England Biolabs) followed by elution of the CDK-as complex after inducible cleavage of the tag in 100 mM DTT at 4 °C.

Purification of the GST-tagged proteins GST-Cla4, GST-Cla4^KD^ (K594A; Tjandra et al., 1998), GST-Kin2 and GST-Kin2^KD^ (D248A; Ghosh et al., 2018) using glutathione beads was carried out as previously described (Geymonat et al., 2009). Resin-bound fractions were stored in 50% v/v glycerol in 1x PBS at -20 °C or proteins eluted with 20 mM reduced glutathione were dialyzed overnight at 4 °C in buffer A and stored.

### Western blot analysis, kinase and binding assays

Expression level of Lte1^WT^-GFP or mutant versions was determined by western blot analysis of whole cell extracts resolved in an 8% SDS-PAGE gel as previously described (Ten Hoopen et al., 2012). GFP-tagged proteins were detected with a mouse monoclonal anti-GFP antibody mix (clones 7.1 and 13.1; Roche, used at 1:1000 dilution) and α-tubulin as loading control with monoclonal antibody B-5-1-2 (Sigma) used at 1:2000 dilution. Myc-tagged proteins were detected using mouse monoclonal 9E10 anti-Myc (Sigma) and His_6_-tagged fusions were detected with anti-His mouse monoclonal antibody (Novagen) at 1:1000 dilution. For detection by chemiluminescence using goat anti-mouse HRP-conjugated secondary antibody (Invitrogen, Fisher Scientific), blots were exposed to film or imaged in an Odyssey XF Imaging System (LI-COR Biosciences) in the chemiluminescence channel. For quantitative analysis the secondary antibody was goat anti-mouse IgG H&L (IRDye 680RD) and the blot was imaged in the 700 nm channel of the Odyssey Imaging System.

Lte1 mobility shift reflecting bulk phosphorylation *in vivo* was assessed by western blot analysis of TCA whole extracts from cells expressing Lte1-ProA fusions at endogenous level. Briefly, early log YEP-dextrose cultures were synchronized in the presence of 15 µg/mL nocodazole for 2.5 h at 25 °C and harvested by centrifugation. After washes in 1 mL cold H_2_O and 1 mL 20% TCA, cells were re-suspended in 200 µL 20% TCA and disrupted with glass beads in a FastPrep-24 grinder followed by addition of 400 µL 5% TCA and a further round of disruption. Lysates were collected and centrifuged for 10 min at 16,000 x g. The resulting pellet was re-suspended in 200 µL 2.5x Laemmli buffer and 50 µL 1M Tris base. Samples were heated at 95 °C for 5 min and subjected to 6% SDS-PAGE followed by western blot analysis. Direct detection of Lte1-ProA constructs was carried out with PAP (peroxidase anti-peroxidase) antibody (Sigma-Aldrich) used at 1:500 dilution.

Coimmunoprecipitation of wild type or mutant Lte1-ProA/His_6_-Ras2 complexes was carried out in a 200 µL reaction containing 3 mg of whole cell extract, 30 µL of rabbit IgG agarose beads in binding buffer (50 mM Tris-HCl pH 7.5, 150 mM NaCl, 1% NP-40, 1 mM DTT, 10% glycerol, Roche cOmplete-EDTA free protease inhibitors, Roche PhosSTOP phosphatase inhibitor cocktail, 2 mM PMSF) incubated at 4 °C for 2 h. After 5 washes with washing buffer (50 mM Tris-HCl pH 7.5, 150 mM NaCl, 1% NP-40, 1 mM DTT), beads were resuspended in 60 µL 2.5 x Laemmli buffer and placed 5 min at 95 °C followed by 10% SDS-PAGE and western blot analysis.

Protein kinase assays with purified components were carried out as follows. The kinase reaction contained 4 to 5 µg of MBP-Lte1 substrates bound to amylose beads, 40 µM ATP, 2 µCi [γ-^32^P] ATP (3000 mCi/mmol; 10mCi/mL, Perkin-Elmer) and 15 to 25 ng of purified kinase in kinase buffer (50 mM Hepes pH 7.5, 10 mM MgCl_2_, 2 mM MnCl_2_ and 1 mM DTT) in a final volume of 30 µL. After a 30 min incubation at 30 °C with gentle tapping to maintain the beads suspended, reactions were stopped by addition of 10 µL of 5x Laemmli buffer and incubation at 95 °C for 3 min. 20 µL aliquots of the phosphorylation reaction were analyzed on an 8% SDS-PAGE gel. The gel was stained with InstantBlue Coomassie solution (Abcam), dried on 3MM blotting paper and exposed for up to 18 hours. The kinases assayed were GST-Cla4, GST-Cla4^KD^, GST-Kin2, GST-Kin2^KD^ and Clb2/Cdc28-as. When indicated, Cdc28-as was inhibited by addition of 5 µM 1NM-PP1 (Geymonat et al., 2020).

*In vitro* pull-down reactions containing ∼10 µg MBP-Lte1 versions bound to amylose resin and 5 µg soluble H-Ras^V12^-myc in 300 µL binding buffer (50 mM Tris-HCl pH 7.5, 150 mM NaCl, 0.2% NP-40, 1 mg/mL BSA, 2 mM DTT) were incubated for 2 h with rotation at 4 °C. After 6 washes with washing buffer (50 mM Tris-HCl pH 7.5, 2 mM DTT, 200 mM NaCl, 0.2% NP-40 for full-length Lte1 baits; 150 mM NaCl, 0.1% NP-40 was used for Lte1^Δ250-950^ baits) the resin was resuspended in 60 µL Laemmli buffer and boiled for 4 min. Pull-down products were resolved on a 10% SDS-PAGE gel. The upper part of the gel (>37kDa) was stained in Coomassie Blue solution for 1 hour and destained overnight to visualize the bait. The lower part of the gel was analyzed by immunoblotting to assess H-Ras^V12^-myc presence using monoclonal 9E10 anti-myc primary antibody. Imaging and quantification were carried out using the Odyssey Imaging System.

For pull-down assays preceded by phosphorylation, 7.5 µg of MBP-Lte1 proteins bound to amylose resin were incubated for 2 h at 37 °C with rotation in a 100 µL reaction containing 1 mM ATP, 50 ng of soluble GST-Kin2 or GST-Kin2^KD^ in kinase buffer (50 mM Hepes pH 7.5, 10 mM MgCl_2_, 2 mM MnCl_2_ and 1 mM DTT). Reactions were stopped by 2 washes of the resin with ice-cold washing buffer. Beads were then resuspended in 300 µL of binding buffer containing 5 µg soluble H-Ras^V12^-myc and incubated with rotation at 4 °C for 2 h. Bound resin was washed 6 times with washing buffer, resuspended in 60 µL Laemmli buffer and boiled for 4 min before resolving by SDS-PAGE as described above.

## Abbreviations

CDK: cyclin-dependent kinase;
CHD: Cdc25 homology domain;
GAP: GTPase-activating protein;
GEF: guanine nucleotide exchange factor;
LRE: Lte1 regulatory element;
MEN: mitotic exit network;
PAK: p21-activated kinase;
REM: Ras exchanger motif;
SPB: spindle pole body;
SPOC: spindle position checkpoint.

## ACKNOWLEDGEMENTS

We thank Olivia Edwards for contributing data to this project. Research reported in this publication was supported, in part, by the Department of Genetics, University of Cambridge (to MS), by a Herchel Smith Scholarship (to SP) and by CSC Cambridge International Scholarships (to QP). The authors declare no competing financial interests.

## SUPPLEMENTARY FIGURE LEGENDS

**Supplementary Figure 1.**
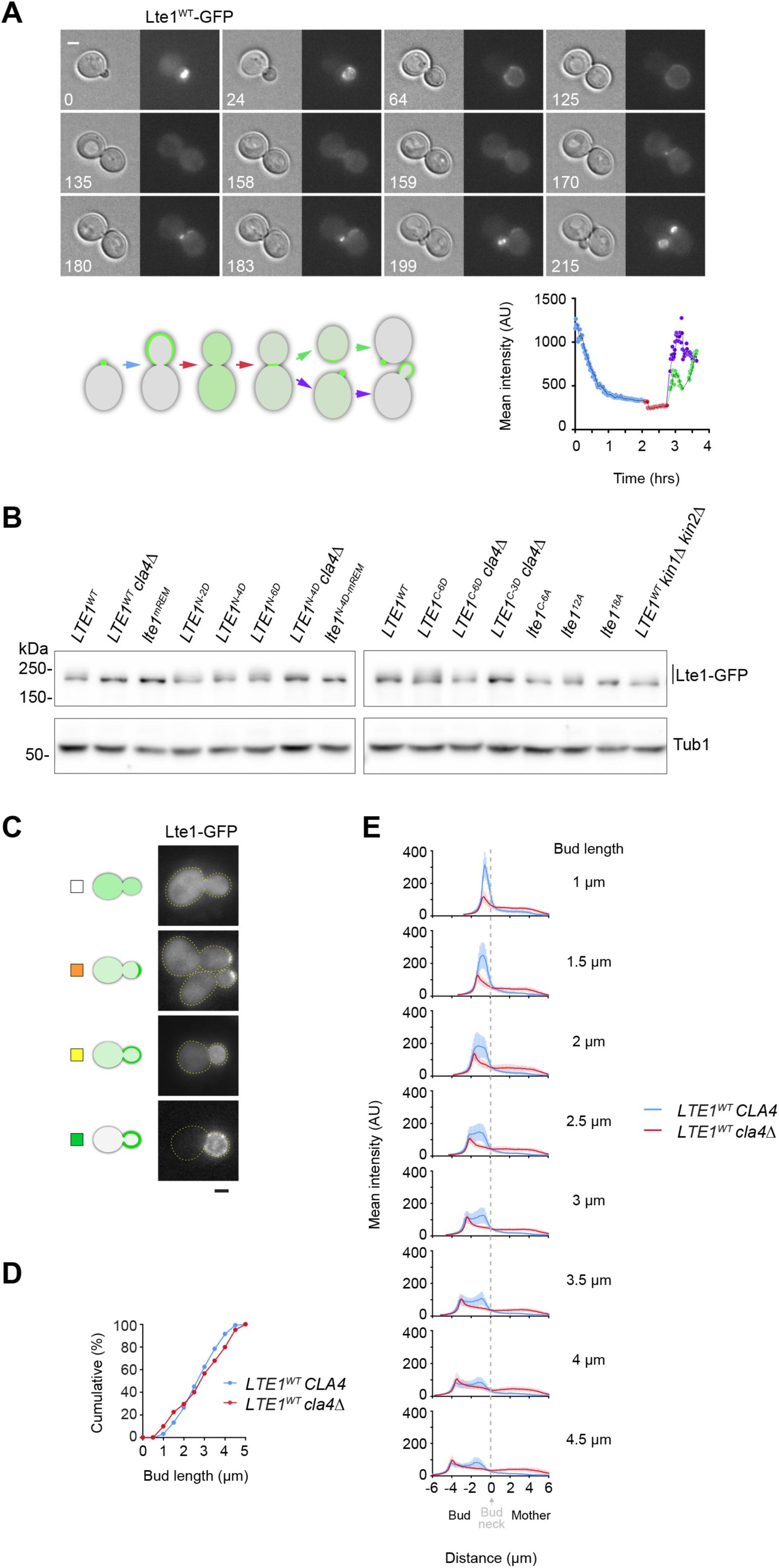
Validating quantitative still imaging analysis of Lte1 cortical localization program. (A) Representative time lapse for Lte1 wild type program of cortical localization at 21 °C. Top panel, selected frames from a time lapse series depicting initial accumulation of Lte1-GFP at the incipient bud (0), progressive cortical spreading during bud growth (24-125), delocalization at mitotic exit with slight localization at the division site (135-159), formation of a label crescent in the daughter cell (159-170) and the initiation of label at a new budding site in the former mother cell (170-onwards) followed later by the accumulation of Lte1 at the new budding site in the former daughter cell (215). Numbers indicate time elapsed in min. Scale bar, 2 µm. Bottom left panel, cartoon depicting the stages of Lte1 cortical localization with colored arrows matching the corresponding section of the plot quantifying mean fluorescence intensity as a function of time elapsed (bottom right panel). Mean fluorescence intensity was maximal shortly after bud emergence and decreased with spreading at the bud cortex during bud growth. This profile is consistent with single cell analysis in cell populations presented in Figure 1 D. (B) Lte1-GFP expression levels assessed by western blot analysis of whole cell extracts of strains used in live imaging studies. Tub1 was used as loading control. (C) Representative modes of Lte1 localization tallied in cell populations in Figures 1 C, 2 C, 2 F, 5 B, 6 F, 7 B and Suppl. Figures 2 B and 3 C. (D) Cumulative distributions of bud length in asynchronous cell populations of wild type vs *cla4Δ* cells quantified in Figure 1 D. (E) Mean fluorescence intensity linescans (solid lines) ± SD (shaded area) compiled from the same dataset analyzed in Figure 1 D, according to bud length (± 0.25 µm). Note that fluorescence in the mother compartment reflects excess background of dispersed Lte1-GFP in the *cla4Δ* mutant.

**Supplementary Figure 2.**
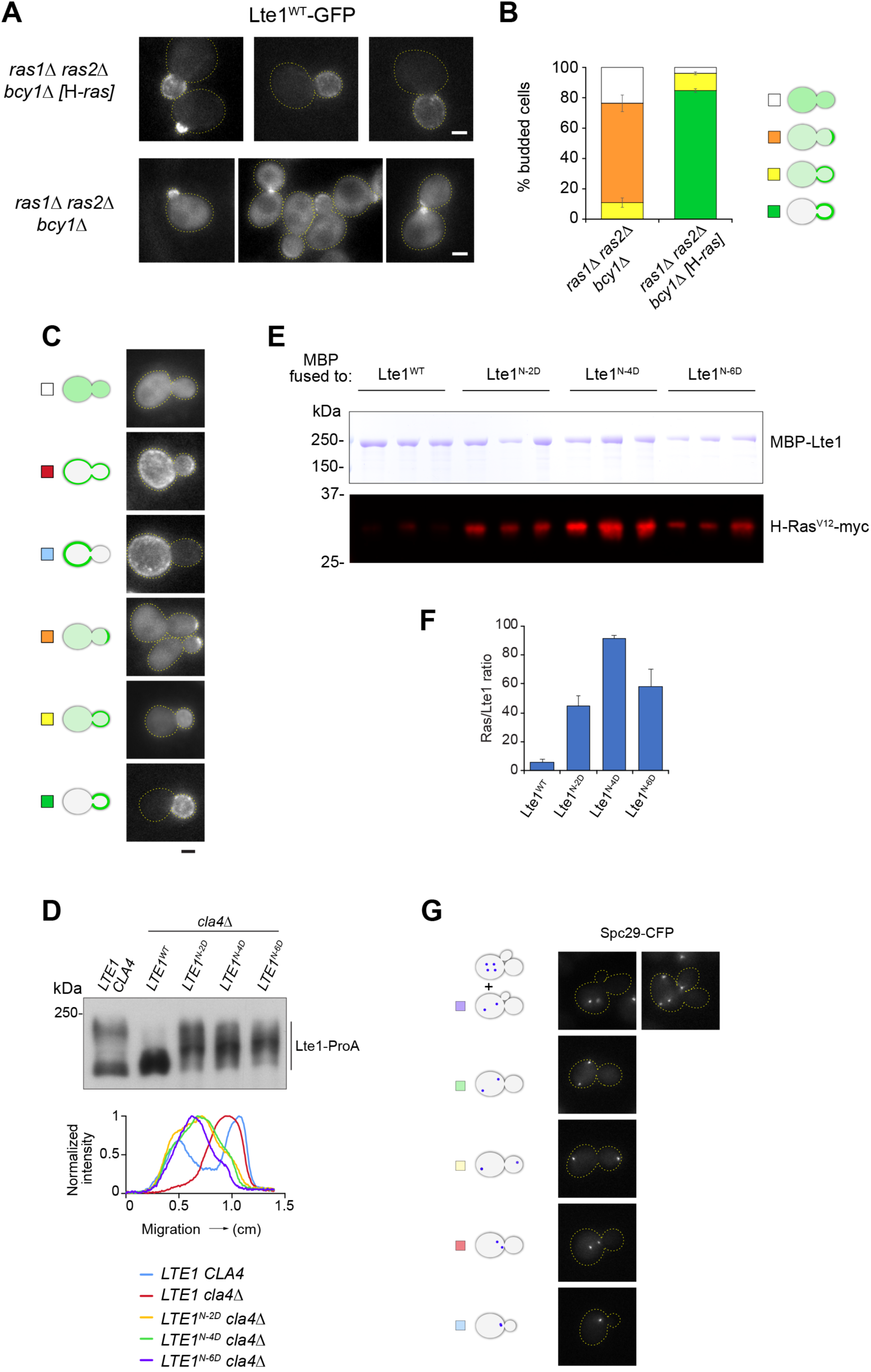
Lte1 anchoring via Ras and the impact of phospho-mimetic substitutions in the N-LRE. (A-B) H-Ras supports Lte1 localization in the absence of yeast Ras proteins. (A) Representative images for Lte1^WT^-GFP localization in r*as1Δ ras2Δ bcy1Δ [*H*-ras]* vs *ras1Δ ras2Δ bcy1Δ* cells. Cells expressing H-Ras efficiently localized Lte1^WT^-GFP. By contrast, cells lacking a Ras protein exhibited limited apical signal over increased cytoplasmic background. Scale bar, 2 µm. N.B. *bcy1Δ* is introduced to suppress *ras1Δ ras2Δ* lethality. (B) Distribution of modes of Lte1^WT^ localization collectively scored in budded cells of the indicated strains. The plot shows the mean ± SEM of three independent counts of 120 cells. (C-G) Impact of phospho-mimetic substitutions at the N-LRE. (C) Representative images for categories of Lte1 cortical localization scored in Figures 3 B, 3 D and 6 B, including those in Suppl. Figure 1 C plus two further classes present in cells carrying phospho-mimetic substitutions at the N-LRE —mother cell cortex mainly (light blue) and mother and bud cell cortex (red). (D) Western blot and linescan analyses along the direction of electrophoretic migration of whole cell extracts of the indicated strains arrested at metaphase. N-LRE phospho-mimetic mutants retained a considerable mobility shift in a *cla4Δ* background. (E-F) Enhanced Lte1 binding to Ras caused by phospho-mimetic substitutions within the N-LRE. (E) After resolving pulled-down products by SDS-PAGE, the top part of the gel was directly stained in Coomassie blue to view MBP-Lte1 baits while the bottom part was subjected to western blot analysis. (F) Mean Ras to Lte1 ratio in triplicate pull-downs ± SD. (G) Representative images for the budded cell categories quantified in Figure 3 H. Unseparated SPBs (light blue), short spindles (red), elongated spindles across the bud neck (yellow), mispositioned anaphase spindles within the mother cell (green), cells undergoing rebudding with mispositioned spindles or further along the subsequent cell cycle following SPB reduplication (violet). Scale bar, 2 µm.

**Supplementary Figure 3.**
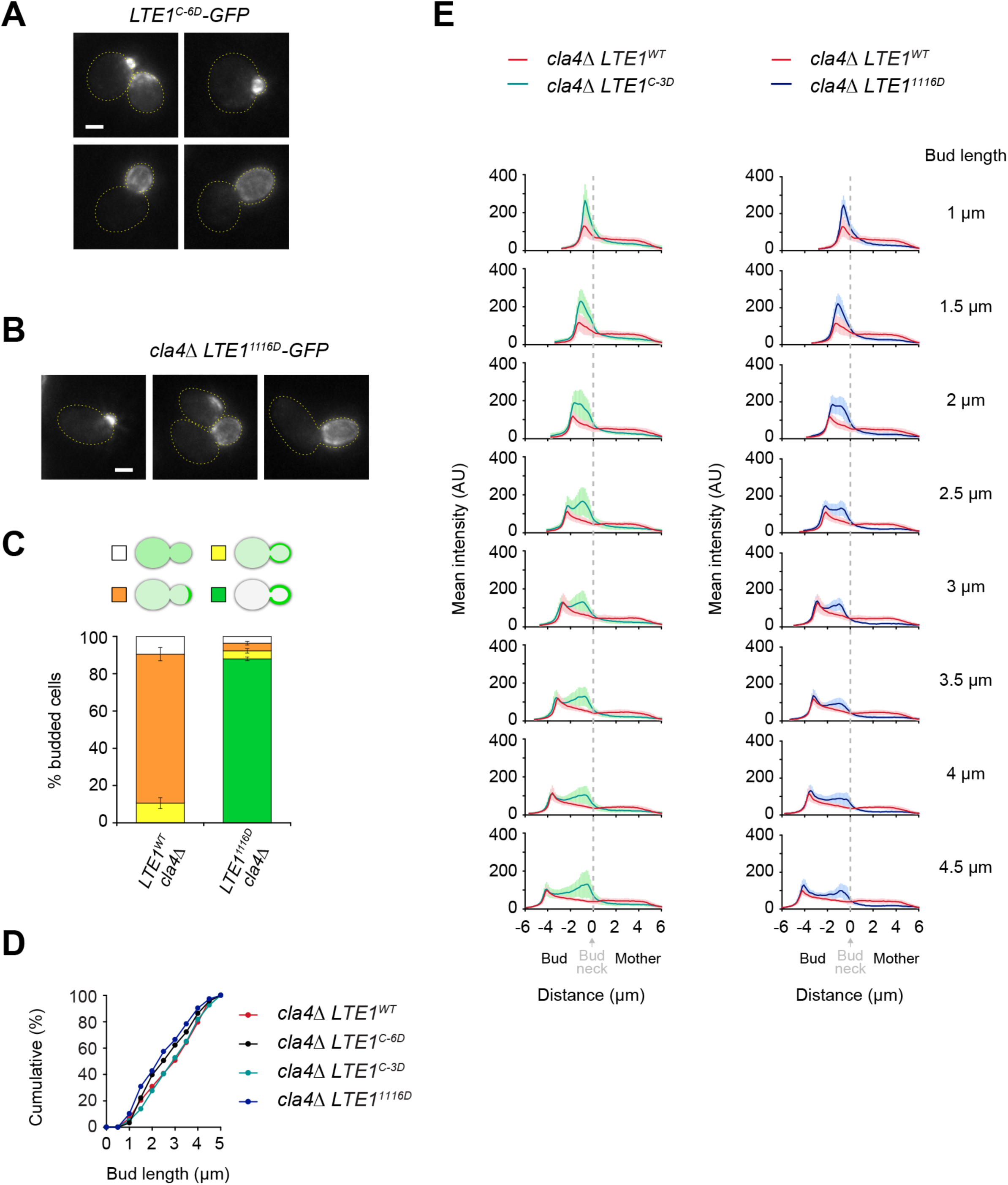
Impact of phospho-mimetic substitutions at the C-LRE on Lte1 localization in *cla4Δ* cells. (A) Phospho-mimetic substitutions at the C-LRE do not perturb Lte1 compartmentalization to the bud. Representative images for Lte1^C-6D^-GFP localization in otherwise wild type cells. Scale bar, 2 µm. (B-C) A single phopho-mimetic substitution at the C-LRE’s PAK site restored Lte1 localization in a *cla4Δ* background. (B) Representative images for localization of Lte1^1116D^ in a *cla4Δ* background. Scale bar, 2 µm. (C) Modes of localization collectively scored in budded cells for Lte1^WT^ vs Lte1^1116D^ expressed in a *cla4Δ* background. The plot shows mean ± SEM of three independent counts of at least 120 cells. (D) Cumulative distributions of bud length in still images of asynchronous cell populations quantified in Figures 5 C and Suppl. Figure 3 E. (E) Mean fluorescence intensity linescans (solid lines) ± SD (shaded area) compiled from *cla4Δ* cells expressing Lte1^WT^-GFP, Lte1^C-3D^-GFP or Lte1^1116D^-GFP, at the indicated bud length (± 0.25 µm). C-3D or 1116D substitutions restored Lte1 localization at the bud cell cortex in a *cla4Δ* mutant although the initial accumulation at the emerging bud did not reach wild type levels (compare with Suppl. Figure 1 E). Also note the suppression of excess cytoplasmic background label within the mother cell compartment relative to that observed for Lte1^WT^ expressed in a *cla4Δ* background.

**Supplementary Figure 4.**
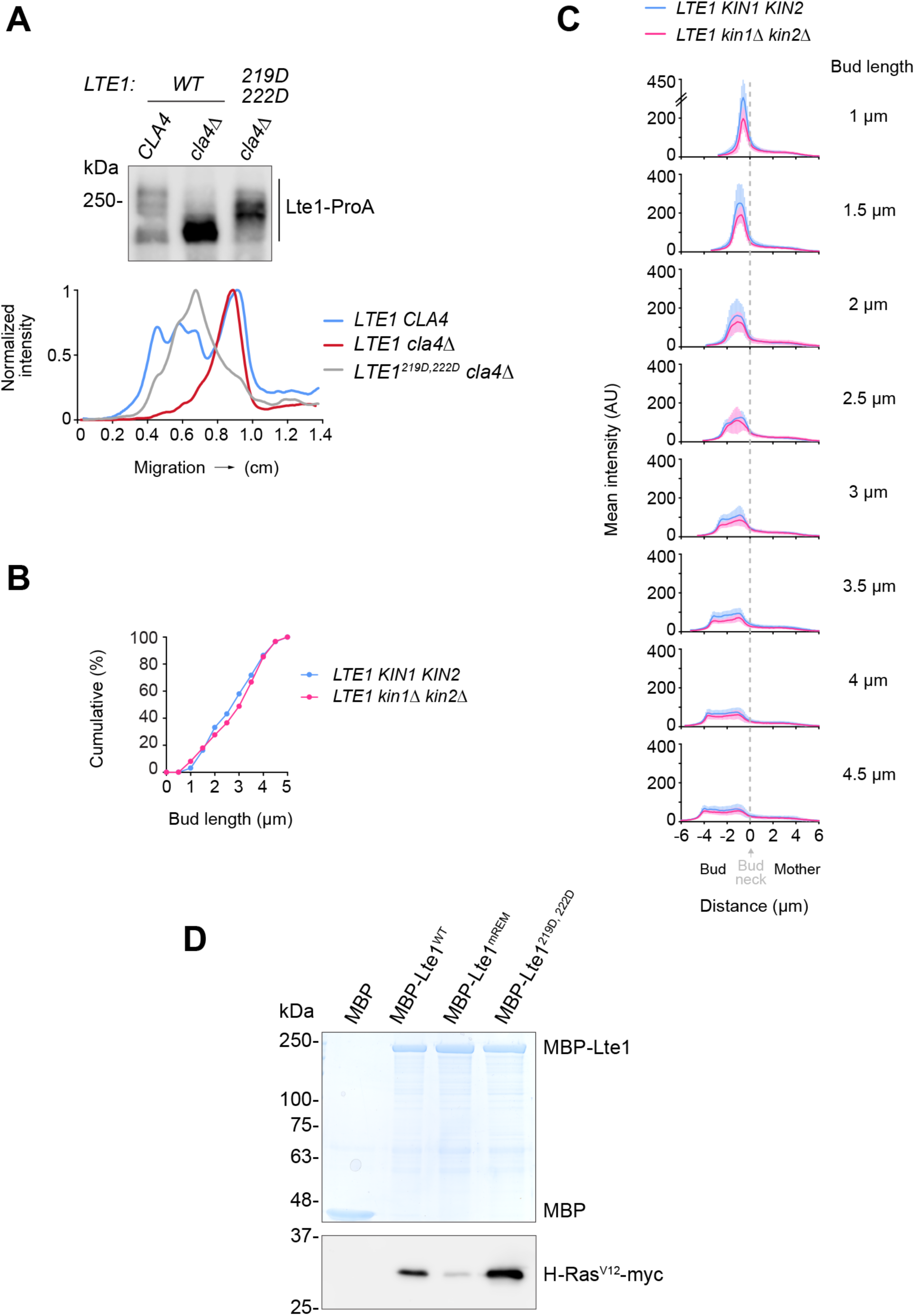
Contribution of Kin1/2 to the Lte1 program of cortical localization. (A) Phospho-mimetic substitutions at positions 219 and 222 restored phospho-dependent mobility shift to Lte1 in a *cla4Δ* background, as assessed by western blot (upper panel) and linescan analyses along the direction of electrophoretic migration (lower panel). (B) Cumulative distributions of bud length in still images of asynchronous cell populations quantified in Figure 6 G and Suppl. Figure 4 C. (C) Mean fluorescence intensity linescans (solid lines) ± SD (shaded area) compiled from cells of the indicated bud length (± 0.25 µm) from the same dataset analyzed in Figure 6 G. Overall Lte1 recruitment to the bud cell cortex was reduced in a *kin1Δ kin2Δ* background. (D) Interaction between Lte1^219D, 222D^ and Ras-GTP assessed by *in vitro* pull down assay. Relative to Lte1^WT^, phospho-mimetic substitutions at positions 219 and 222 enhanced binding to Ras-GTP. In agreement with Figure 3 F, MBP-Lte1^mREM^ exhibited impaired binding.

**Supplementary Figure 5.**
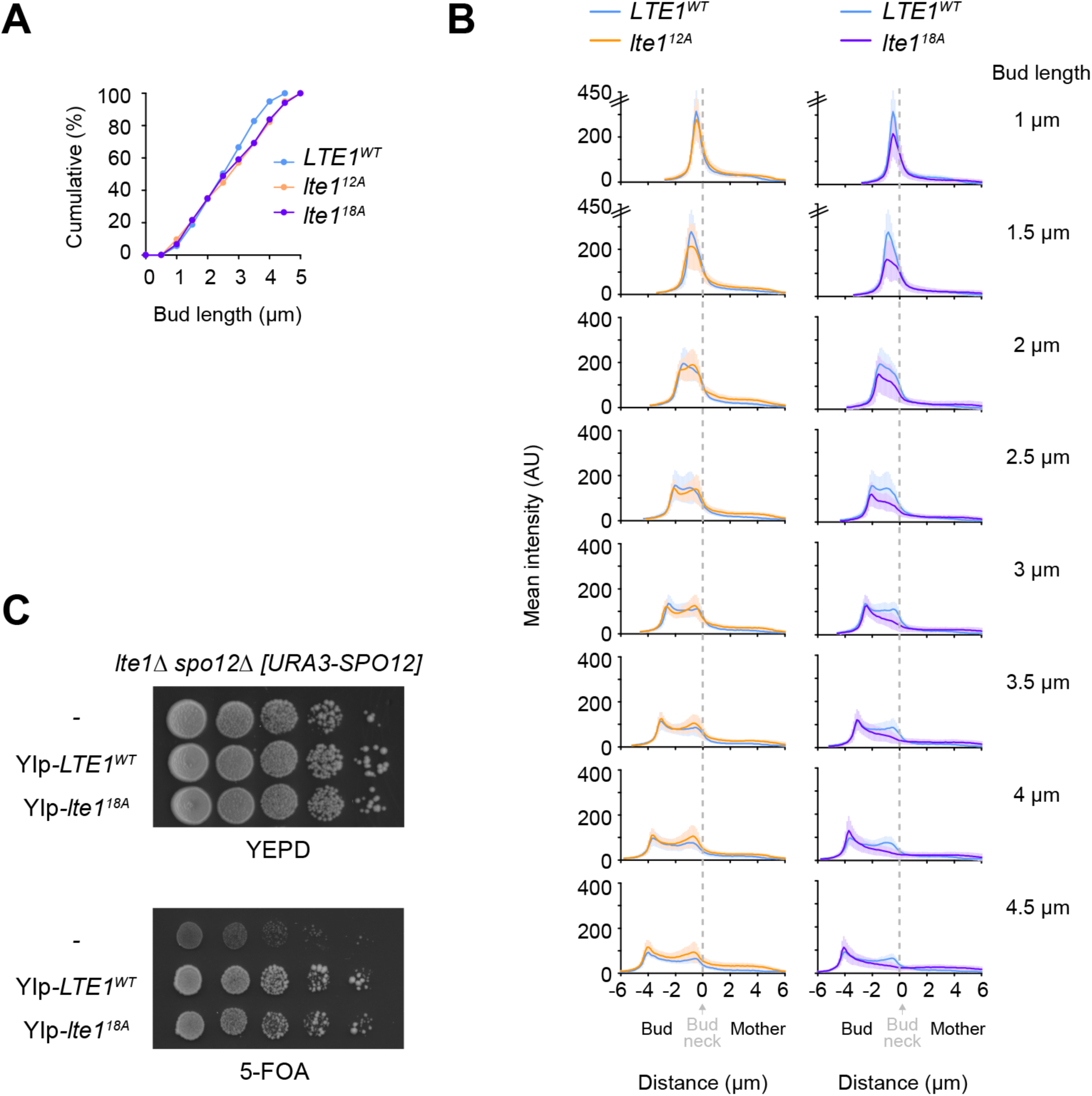
**Further characterization of *lte1^18A^*** (A) Cumulative distributions of bud length in the cell populations analyzed in Figure 7 C. (B) Mean fluorescence intensity linescans (solid lines) ± SD (shaded area) compiled from cells of the indicated bud length (± 0.25 µm) from the same dataset analyzed in Figure 7 C. (C) *lte1^18A^* can suppress *lte1Δ spo12Δ* co-lethality as observed by the ability of transformants to grow upon counter-selection of the resident *URA3-SPO12* plasmid on 5-FOA medium.

---

**Supplementary Table 1.**
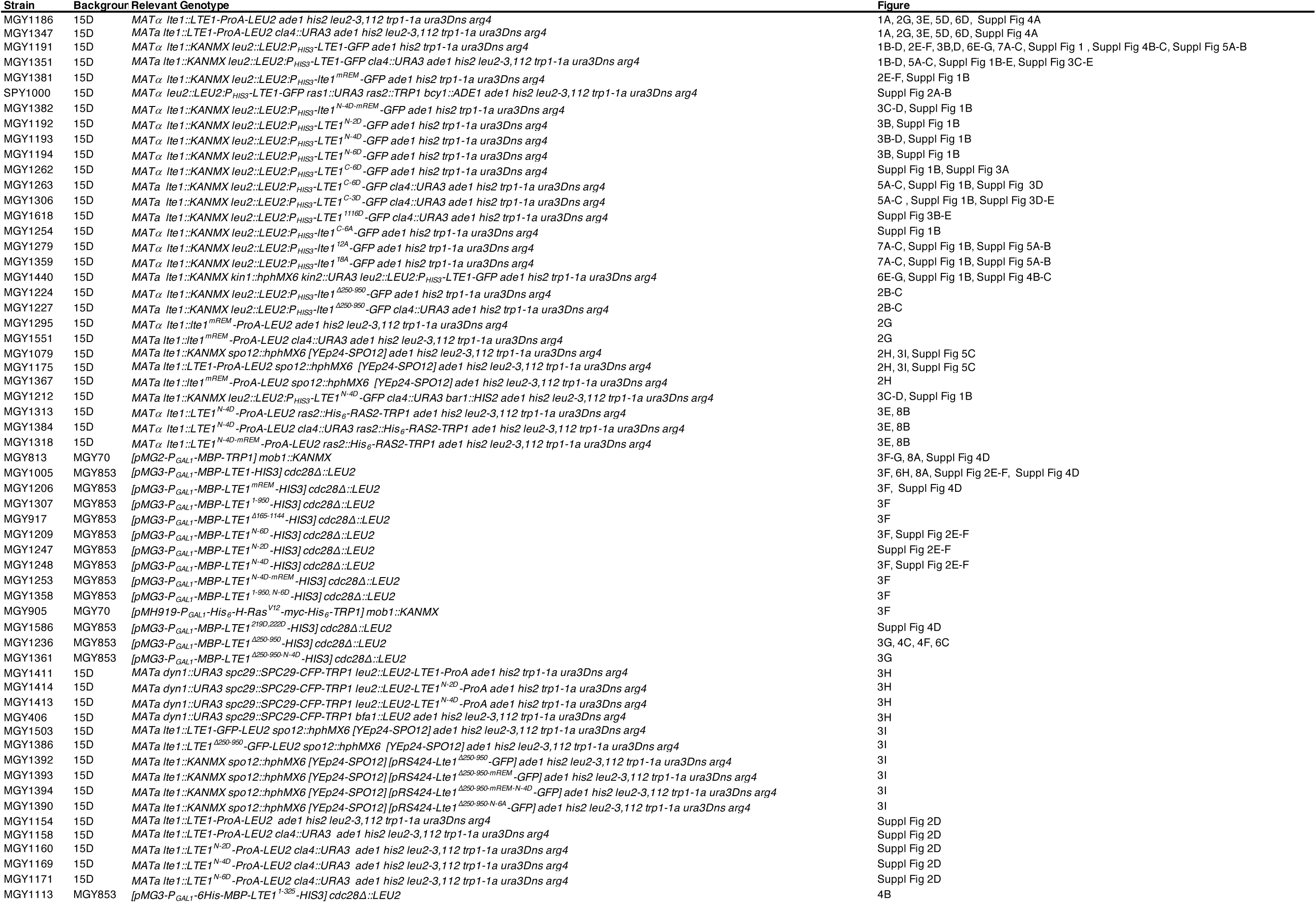
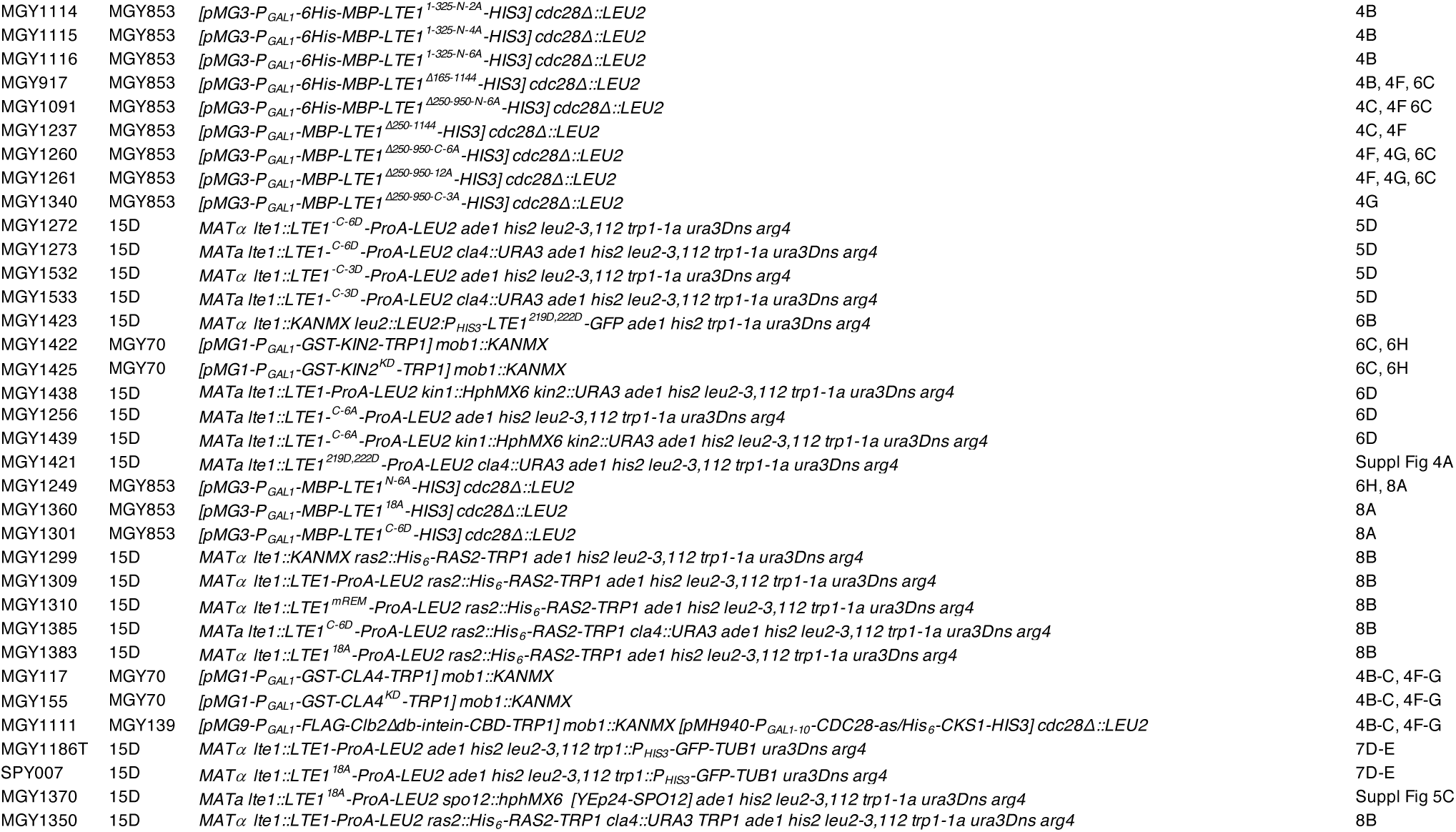
Yeast strains used in this study.

